# HERC3 E3 ligase provides an ERAD branch eliminating select membrane proteins

**DOI:** 10.1101/2023.10.16.562477

**Authors:** Yuka Kamada, Yuko Ohnishi, Chikako Nakashima, Aika Fujii, Mana Terakawa, Ikuto Hamano, Uta Nakayamada, Saori Katoh, Noriaki Hirata, Hazuki Tateishi, Ryosuke Fukuda, Hirotaka Takahashi, Gergely L. Lukacs, Tsukasa Okiyoneda

## Abstract

Aberrant proteins located in the endoplasmic reticulum (ER) undergo rapid ubiquitination by multiple ubiquitin (Ub) E3 ligases and are retrotranslocated to the cytosol as part of the ER-associated degradation (ERAD). Despite several ERAD branches involving different Ub E3 ligases, each with distinct substrate specificity, the molecular machinery responsible for these ERAD branches in mammalian cells remains not fully understood. In this study, we have discovered a cytosolic Ub ligase called HERC3, which fulfills a distinct role in facilitating the ERAD of select polytopic membrane proteins. Using a series of multiplex knockdown/knockout experiments, we have demonstrated that HERC3 functions independently of the ER-embedded ubiquitin ligases RNF5 and RNF185 (RNF5/185) to facilitate the ubiquitination, retrotranslocation, and ERAD of misfolded CFTR. Furthermore, HERC3 collaborates with RNF5/185 to enhance the association of UBQLN proteins, thereby augmenting the retrotranslocation and ERAD of misfolded CFTR. While RNF5/185 participates in the ERAD process of both misfolded ABCB1 and CFTR, HERC3 specifically promotes the ERAD of CFTR, likely due to its ability to interact with the less hydrophobic membrane-spanning domains of CFTR. HERC3 may detect exposed transmembrane domains on the cytoplasmic surface of the ER, thereby facilitating the recruitment of UBQLN and subsequently accelerating the ERAD of select polytopic membrane proteins.

## Introduction

Conformationally defective proteins in the endoplasmic reticulum (ER) resulting from genetic mutations and environmental stresses are selectively recognized by the ER quality control (ERQC) machinery and eliminated through ER-associated degradation (ERAD). Aberrant ER luminal and membrane proteins undergo retrotranslocation from the ER to the cytoplasm and ubiquitination, both of which are coordinated by the ubiquitin (Ub) ligase complexes (Christianson and Ye, 2014; Ruggiano et al., 2014). In mammalian cells, ERAD pathways involve at least 10 Ub ligases, each defining specific ERAD branches with specificity towards certain classes of substrates (Krshnan et al., 2022). Certain Ub ligases work together in concert to promote ERAD. For instance, at the ER membrane, Gp78 (AMFR) elongates the Ub chains initiated by RNF5 (RMA1) to facilitate ERAD (Morito et al., 2008). Additionally, cytoplasmic HECT Ub ligase UBE3C collaborates with the ER-embedded Ub ligase RNF185/MBRL complex to promote ERAD (van de Weijer et al., 2020). These various ERAD branches ultimately converge on the cytosolic p97/VCP complex, which extracts ubiquitinated substrates from the ER membrane for proteasomal degradation (Wu and Rapoport, 2018). Several Ub ligases, including Hrd1 (Schoebel et al., 2017; Vasic et al., 2020; Wu et al., 2020) and Doa10 (Schmidt et al., 2020), also appear to function as the retrotranslocation channel. Cytoplasmic chaperones including Bag6 (Wang et al., 2011; Xu et al., 2012) and proteasome shuttling factor UBQLNs (Lim et al., 2009) function downstream of p97-mediated substrate extraction to promote the delivery of ubiquitinated substrates to the proteasome. While the ERQC mechanism is necessary to maintain cellular proteostasis and physiological function, it is also involved in the pathogenesis of diseases such as cystic fibrosis (CF) which is caused by mutations of cystic fibrosis transmembrane conductance regulator (CFTR).

CFTR is a 1480-residue polytopic membrane glycoprotein that is predicted to contain two membrane-spanning domains (MSD) with six transmembrane segments, two large cytosolic nucleotide-binding domains (NBD), and a cytosolic regulatory (R) domain (Riordan, 2008; Riordan et al., 1989). It belongs to the ATP-binding cassette (ABC) transporter family and functions as a cAMP-regulated Cl^-^ channel at the apical plasma membrane (PM) of epithelial cells and its mutations cause CF, one of the most common genetic diseases in Caucasians (Boucher, 2004; Riordan, 2008; Riordan et al., 1989). The most common mutation in CF is ΔF508-CFTR in which phenylalanine at position 508 is deleted in the NBD1 located in the cytoplasmic region (Rich et al., 1990; White et al., 1990). The ΔF508 mutation destabilizes NBD1 and the interdomain assembly of CFTR (Du et al., 2005; He et al., 2013; Lewis et al., 2005; Rabeh et al., 2012). Consequently, misfolded ΔF508-CFTR is ubiquitinated and prematurely degraded by the proteasome, resulting in marginal cell surface expression (Jensen et al., 1995; Ward et al., 1995). Several Ub E3 ligases have been identified for CFTR ubiquitination at the ER. A chaperone-associated cytosolic E3 ligase CHIP (STUB1) (Meacham et al., 2001), ER-embedded E3 ligases RNF5 (Younger et al., 2006), RNF185 (El Khouri et al., 2013), and Gp78 (Morito et al., 2008) are involved in CFTR ubiquitination possibly at the distinct biosynthesis stages and at multiple sites within CFTR polypeptides (Oberdorf et al., 2006). Cytoplasmic Ub ligase CHIP appears to recognize the conformational defects of cytoplasmic regions of CFTR including NBD1 coordinated with the Hsc70/Hsp70 and Hsp90 chaperone complex (Younger et al., 2006; Younger et al., 2004). ER-embedded RNF5 appears to recognize the N-terminal regions of CFTR, such as MSD1 (Younger et al., 2006), and facilitates polyubiquitination in cooperation with Gp78 (Morito et al., 2008). RNF185, a paralog of RNF5, may recognize the CFTR’s MSDs in coordination with MBRL and TMUB1/2 based on a previous study (van de Weijer et al., 2020) although the exact recognition mechanism remains unknown. The ubiquitinated CFTR undergoes retrotranslocation which may be mediated by Derlin-1 (Sun et al., 2006; Wang et al., 2008) and p97 complex (Carlson et al., 2006). Derlin-1 is thought to promote the extraction of MSD1 in the ubiquitinated CFTR (Sun et al., 2006). The extraction of the transmembrane segments is considered generally rate-limiting in the degradation of polytopic membrane proteins. The p97 complex could specifically extract the transmembrane segments of CFTR to accelerate ERAD (Carlson et al., 2006). The multiple Ub ligases appear to recognize various features of CFTR’s conformational defects located in multiple regions and utilize distinct downstream ERAD branches to efficiently eliminate misfolded CFTR in the early secretory pathway. However, due to a lack of comprehensive analysis, the CFTR ERAD branches have not been fully understood. Moreover, how multiple Ub ligases, including ER-embedded and cytoplasmic E3 ligases, coordinately regulate ERAD processes, including ubiquitination and retrotranslocation, still requires further investigation.

In this study, we have identified a cytoplasmic Ub ligase HERC3 that plays a crucial role in a novel ERAD branch dedicated to the selective degradation of misfolded CFTR. Through our HiBiT-based ERAD and retrotranslocation assays, we have demonstrated that both ER-embedded RNF5/185 and cytosolic HERC3 independently promote the retrotranslocation and ERAD of misfolded CFTR. Specifically, HERC3 facilitates the association of misfolded CFTR with UBQLN2, which in turn promotes retrotranslocation. Unlike RNF5/185, HERC3 displays selectivity for certain membrane proteins, such as CFTR, possibly by interacting with the MSDs. Our results suggest that cytoplasmic HERC3 may function as an ERQC-associated E3 ligase that interacts with the transmembrane segments exposed on the surface of the ER membrane, thereby providing an ERAD branch specialized for a specific type of membrane proteins.

## Result

### HERC3 limits the cell surface expression of ΔF508-CFTR by facilitating the ERAD

Previously we conducted a comprehensive siRNA screening in CFBE cells stably expressing ΔF508-CFTR-HRP and identified RFFL Ub ligase whose knockdown (KD) increased the PM level of rescued (r)ΔF508-CFTR which was forcibly expressed at the PM by low-temperature (26°C) incubation (Okiyoneda et al., 2018). At the same time, we have identified a novel Ub ligase HERC3 whose KD also increased the rΔF508-CFTR PM level (Figure 1A). We ruled out the possibility of this effect being due to an off-target effect, as the increased PM CFTR was also observed in HERC3 KD using individual siRNAs (Figure 1A). The efficacy of these siRNAs was confirmed through reverse transcription quantitative PCR (RT-qPCR) analysis (Figure 1B). To determine if the increased PM CFTR was functional upon HERC3 KD, we performed a halide-sensitive YFP quenching assay in CFBE Teton cells (Okiyoneda et al., 2018). As expected, the YFP quenching induced by CFTR-mediated iodide influx was increased in the cells transfected with siHERC3, indicating that HERC3 KD significantly increased the functional rΔF508-CFTR channel at the PM (Figure 1C). Further analysis through Western blotting revealed that HERC3 KD increased both mature and immature rΔF508-CFTR after low-temperature rescue (Denning et al., 1992), but it increased the immature ΔF508-CFTR at 37°C, suggesting that HERC3 might regulate the CFTR level at the ER (Figures 1D and 1E). To investigate the impact of HERC3 KD on CFTR ERAD, we conducted a cycloheximide (CHX) chase experiment. The cell-based ELISA showed that HERC3 KD slightly but significantly decelerated the elimination of immature ΔF508-CFTR in CFBE cells (Figure 1F). However, HERC3 KD did not influence the PM stability of rΔF508-CFTR, in contrast to the effect observed with KD of RFFL which is involved in the elimination of rΔF508-CFTR at the PM and endosomes (Okiyoneda et al., 2018) (Figure 1G). These findings suggest that the increased level of mature rΔF508-CFTR is not due to the reduced peripheral degradation of mature CFTR but could be attributed to the decelerated ERAD of immature CFTR. In line with this hypothesis, HERC3 KD resulted in reduced ubiquitination of immature ΔF508-CFTR-3HA, which is fused with an N-terminal histidine-biotin-histidine (HBH) tag (HBH-ΔF508-CFTR) (Okiyoneda et al., 2018; Tagwerker et al., 2006) (Figure 1H). Conversely, when HERC3 was overexpressed, it led to a decrease in immature ΔF508-CFTR levels, and this effect depended on the catalytic activity of the HECT domain as the deletion or catalytically inactive C1018A mutation of the HECT domain abolished this effect (Figures 1I and 1J). Furthermore, deleting the RCC1 Like Domain (RLD) of HERC3 also reduced its interaction with ΔF508-CFTR and the subsequent decrease in immature ΔF508-CFTR expression (Figures 1J and 1K), suggesting that the RLD is essential for HERC3’s interaction with immature ΔF508-CFTR at the ER.

**Figure 1.**
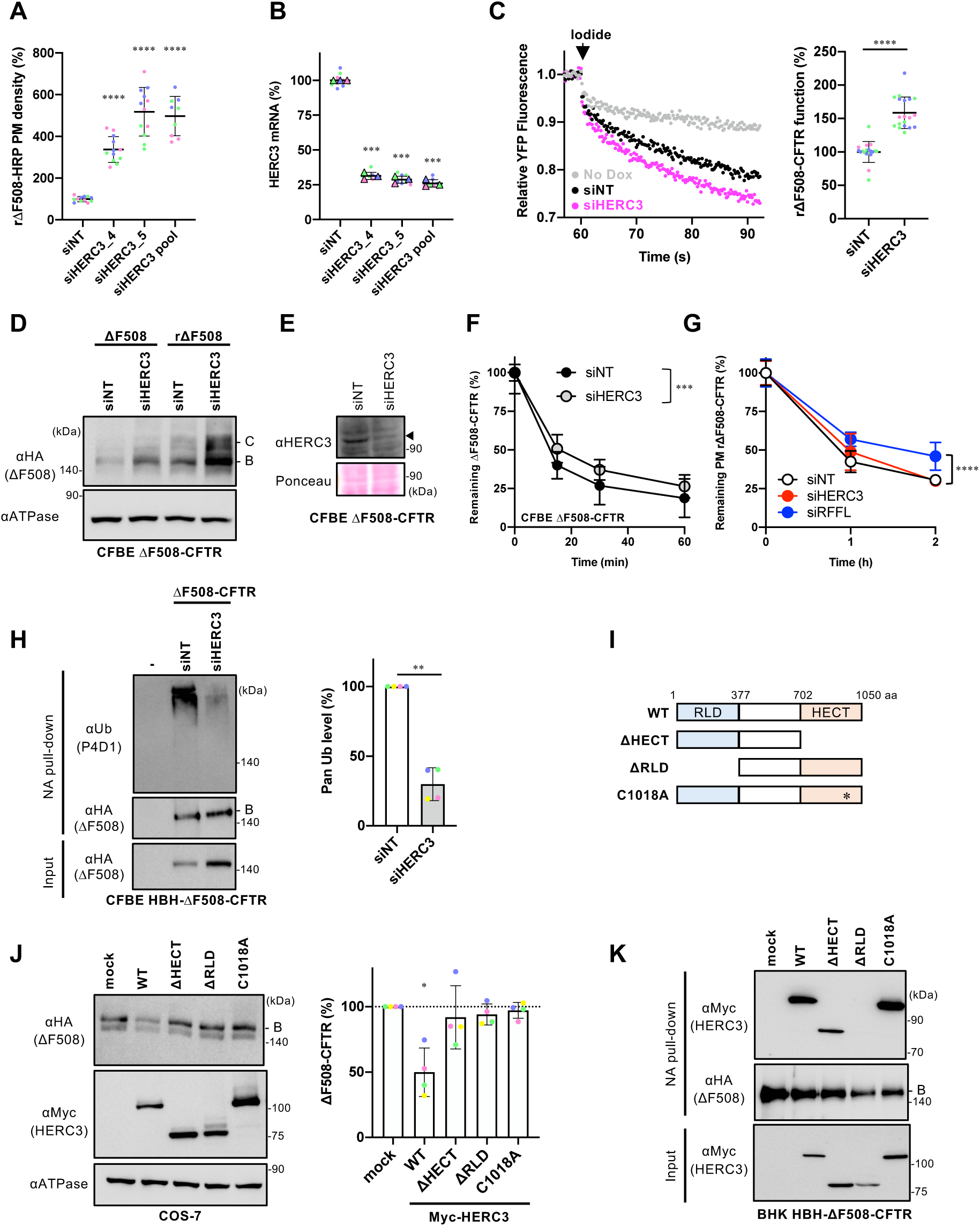
HERC3 participates in the ubiquitination and ERAD of ΔF508-CFTR. (A) The PM density of rΔF508-CFTR-HRP in CFBE Teton cells transfected with 50 nM siNT or siHERC3, as indicated (n=9-12). Each independent experiment, consisting of 3-4 biological replicates (n), is color-coded. (B) Quantitative PCR analysis assessed HERC3 KD efficiency in CFBE Teton ΔF508-CFTR-HRP cells (n=3). Each biological replicate (n) is color-coded: the averages from 3 technical replicates are shown in triangles. (C) The channel function of rΔF508-CFTR-3HA in CFBE Teton cells transfected with 50 nM siNT or siHERC3 pool was measured by YFP quenching assay. The initial YFP quenching rate was quantified as the CFTR function (right, n=19). Each independent experiment, consisting of 4-8 biological replicates (n), is color-coded. (D) Western blotting analyzed steady-state levels of ΔF508-CFTR-3HA with (rΔF508) or without 26°C rescue (ΔF508) in CFBE Teton cells transfected with 50 nM siNT or siHERC3 pool. Na^+^/K^+^ ATPase (ATPase) was used as a loading control. B, immature form; C, mature form. (E) Western blotting confirmed HERC3 KD in CFBE Teton ΔF508-CFTR-3HA cells. Ponceau staining was used as a loading control. A filled triangle indicates HERC3. (F) Cellular ΔF508-CFTR-3HA stability in CFBE Teton cells transfected with 50 nM siNT or siHERC3 pool was measured by cell-based ELISA using an anti-HA antibody after CHX treatment (n=12). (G) The PM stability of rΔF508-CFTR-3HA in CFBE cells transfected with 50 nM siNT, siRFFL pool, or siHERC3 pool was measured by PM ELISA (n=12 biological replicates). (H) Ubiquitination levels of HBH-ΔF508-CFTR-3HA in CFBE Teton cells were measured by Neutravidin (NA) pull-down under denaturing conditions (NA pull-down) and Western blotting. The CFTR ubiquitination level was quantified by densitometry and normalized to CFTR in precipitates (right, n=4). (I) A schematic diagram of the HERC3 domain composition with the residue numbers at the domain boundaries. HERC3 mutants used in this study are also shown. (J) The effects of overexpressed Myc-HERC3 variants on the steady-state level of ΔF508-CFTR-3HA were analyzed by Western blotting in co-transfected COS-7 cells. The immature ΔF508-CFTR (B band) was quantified by densitometry (right, n=4). (K) The interaction of Myc-HERC3 variants with HBH-ΔF508-CFTR-3HA in BHK cells was analyzed by NA pull-down and Western blotting. ΔF508-CFTR was rescued at 26°C incubation for 2 days, followed by a 1-hour incubation at 37°C (A-C, D, G). Statistical significance was assessed by one-way ANOVA (A) or one-way repeated-measures (RM) ANOVA (B, J) with Dunnett’s multiple comparison tests, a two-tailed unpaired (C) or paired Student’s t-test (H), or two-way ANOVA (F, G). Data represent mean ± SD. *p < 0.05, **p < 0.01, ***p < 0.001, ****p < 0.0001, ns, not significant.

### HERC3 and RNF5/185 additively facilitate ΔF508-CFTR ERAD

Our findings suggest that HERC3 may play a role in facilitating CFTR ERAD, alongside other Ub ligases such as CHIP, RNF5, RNF185, and Gp78. To compare the impact of these Ub ligases on ΔF508-CFTR, we evaluated the effect of individually knocking down each E3 ligase in CFBE cells. Surprisingly, while HERC3 KD significantly increased cellular ΔF508-CFTR levels, KD of CHIP, RNF5, RNF185, or Gp78 had no significant effect (Figure 2A). Upon low-temperature rescue, the cell surface levels of rΔF508-CFTR were increased by KD of HERC3, RNF5, or RNF185, indicating their crucial role in limiting ΔF508-CFTR abundance in CFBE cells (Figure 2B). To examine whether HERC3 collaborates with other CFTR-related E3 Ub ligases, which may have redundant functions, we tested the effect of simultaneously knocking down HERC3 and one of these E3 ligases. While KD of either CHIP or Gp78 did not enhance the effect of HERC3 KD on the PM levels of rΔF508-CFTR, KD of RNF5 or RNF185 significantly enhanced the effect of HERC3 KD (Figure 2B). The absence of an impact from CHIP KD in the CFTR QC in CFBE cells was also documented previously (Okiyoneda et al., 2018), suggesting that the influence of CHIP may fluctuate depending on the cell type. This result suggests that HERC3 may regulate ΔF508-CFTR through a pathway distinct from RNF5 and RNF185.

**Figure 2.**
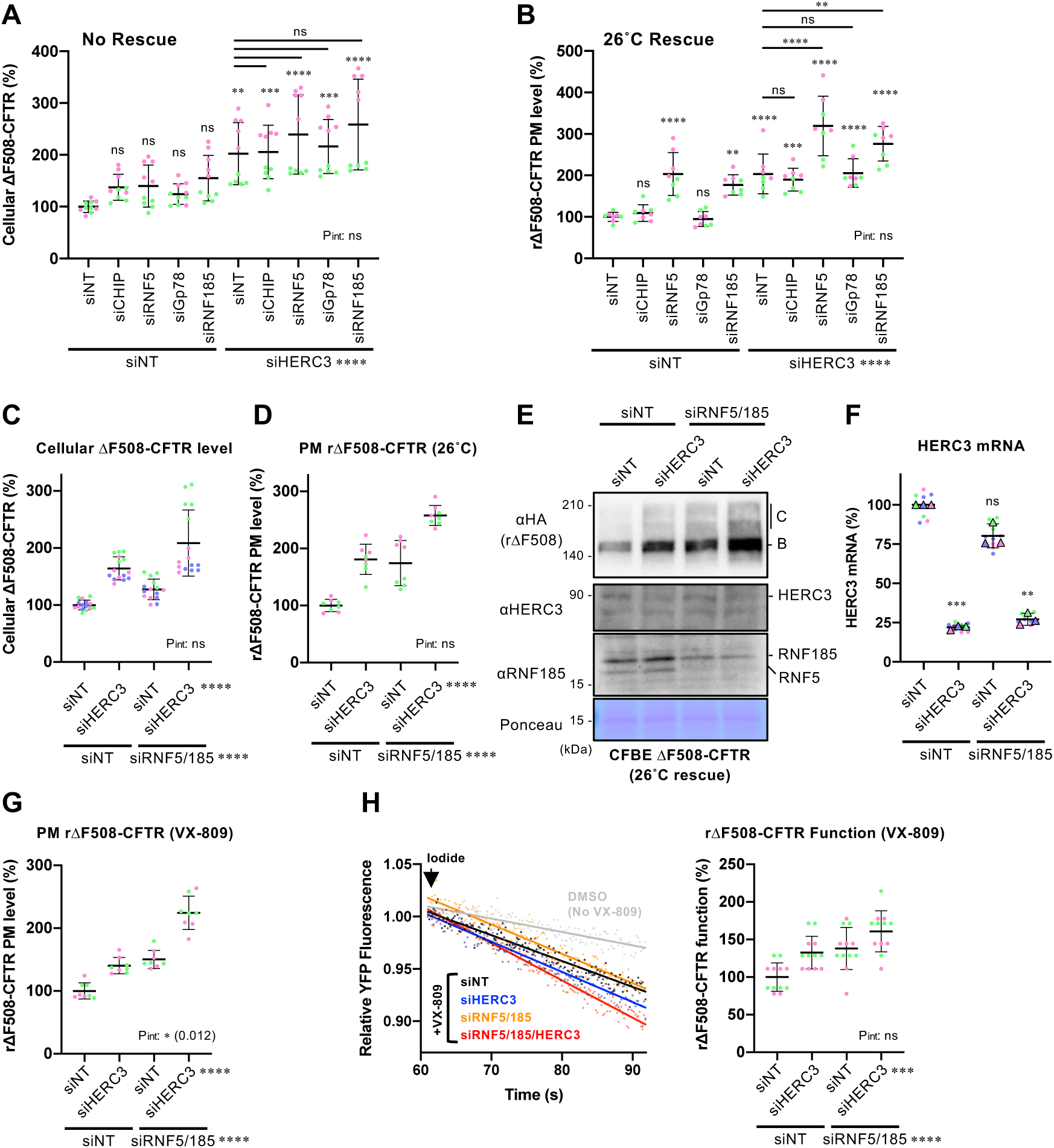
HERC3 and RNF5/185 additively reduce ΔF508-CFTR. (A, B) The cellular level of ΔF508-CFTR-3HA (A, n=10) and PM level of rΔF508-CFTR-HRP (B, n=8) in CFBE Teton cells transfected with 50 nM siRNA indicated was measured by cell-based ELISA using an anti-HA antibody and HRP assay, respectively. (C, D) The cellular level of ΔF508-CFTR-3HA (C, n=15) and PM levels of rΔF508-CFTR-HRP induced by 26°C rescue (D, n=8) in CFBE Teton cells transfected with 50 nM siRNA were measured by ELISA using an anti-HA antibody (C) and HRP assay (D), respectively. (E, F) Western blotting analyzed steady-state levels of rΔF508-CFTR-3HA in CFBE Teton cells transfected with 50 nM siRNA indicated (E). Ponceau staining was used as a loading control. B, immature form; C, mature form. The anti-RNF185 antibody detected both RNF5 and RNF185 because of the cross-reactivity. HERC3 KD was confirmed by quantitative PCR (F, n=3). Each biological replicate (n) is color-coded: the averages from 3 technical replicates are shown in triangles. (G) The PM levels of rΔF508-CFTR-HRP induced by 3 µM VX-809 treatment at 37°C for 24 hours in CFBE Teton cells transfected with 50 nM siRNA indicated (n=8). (H) Representative traces (left) of the YFP fluorescence and quantification of the initial YFP quenching rate (right, n=12) as a measure of rΔF508-CFTR function in CFBE cells transfected with 50 nM siRNA, as indicated. Each independent experiment consisting of 4 (B, D, G), 5 (A, C), or 6 (H) biological replicates (n) is color-coded. Statistical significance was assessed by one-way RM ANOVA with Dunnett’s multiple comparison tests (F) or two-way ANOVA with Holm-Sidak multiple comparison tests which revealed a significant main effect of HERC3 KD or RNF5/185 DKD, but no interaction between them (P_int_>0.05, C, D, H) except for G (P_int_=0.012). Data represent mean ± SD. **p < 0.01, ***p < 0.001, ****p < 0.0001, ns, not significant.

To further investigate this possibility, we assessed the effect of HERC3 KD on ΔF508-CFTR levels upon simultaneous KD of RNF5 and its paralog RNF185 (RNF5/185 DKD), both of which could be functionally redundant in CFTR ERAD (El Khouri et al., 2013). As expected, the impact of HERC3 KD on cellular ΔF508-CFTR levels remained unaffected by siRNF5/185 (Figure 2C). Consequently, the combined KD of HERC3 and RNF5/185 resulted in an additive increase in cellular CFTR levels (Figure 2C). This finding suggests that HERC3’s regulation of ΔF508-CFTR operates independently of RNF5 and RNF185. In line with these findings, the simultaneous KD of HERC3 and RNF5/185 led to an additive increase in the PM level of low-temperature rescued ΔF508-CFTR (Figure 2D). Similarly, comparable results were observed for the overall CFTR protein levels (Figures 2E and 2F). HERC3 KD also increased the PM levels of rΔF508-CFTR induced by the CFTR corrector VX-809 treatment in CFBE cells, and this effect was further enhanced upon RNF5/185 DKD (Figure 2G). VX-809 has been reported to partially correct the CFTR conformational defects such as defective MSD1/2-NBD1 interaction by binding with its MSD1, inducing partial PM expression (Farinha et al., 2013; Fiedorczuk and Chen, 2022a; Loo et al., 2013; Okiyoneda et al., 2013; Ren et al., 2013; Van Goor et al., 2011). Additionally, KD of both HERC3 and RNF5/185 additively increased the functional ΔF508-CFTR channel at the PM in the presence of VX-809 (Figure 2H). These findings suggest that, in addition to the RNF5/185 pathway, the HERC3 ERAD branch limits the efficacy of the CFTR corrector. Combining strategies to counteract both ERAD branches may enhance the effectiveness of CFTR correctors.

To accurately assess the role of HERC3 in CFTR ERAD, we developed a live cell CFTR degradation assay that allowed us to measure the degradation kinetics of ΔF508-CFTR fused with the HiBiT tag in the C-terminal cytosolic region (ΔF508-CFTR-HiBiT(CT)) in real-time (Figure 3A). In this assay, we measured the NanoLuc (Nluc) luminescence signal induced by the association of the HiBiT tag with the co-expressed LgBiT in 293MSR cells at 37°C. The luminescence signal gradually attenuated during the CHX chase, and its half-life (t_1/2_) was approximately 70 minutes (Figure 3B). The addition of a proteasome inhibitor MG-132 significantly reduced the luminescence attenuation during the CHX chase, indicating that the luminescence attenuation represents the kinetic proteasomal degradation of ΔF508-CFTR-HiBiT(CT) (Figure 3B). The degradation kinetics of ΔF508-CFTR in the HiBiT-based assay closely resembled those observed in traditional Western blot analyses (Figure 3C), validating the reliability of the new HiBiT degradation assay. However, in Western blot analyses, the ERAD rate seemed significantly faster, potentially because of its lower sensitivity in detecting CFTR. Using this innovative assay, we examined the effect of HERC3 KD on ΔF508-CFTR ERAD in 293MSR wild-type (WT) cells and in cells with the double knockout (KO) of RNF5 and RNF185 (5/185 DKO) that we established using the CRISPR-Cas9 system (Figures S1A and S1B). Western blotting and RT-qPCR confirmed the RNF5/185 DKO and HERC3 KD in 293MSR cells (Figures 3D and 3E). The live cell degradation assay showed that consistent with the results in CFBE cells, HERC3 KD modestly decelerated the ERAD of ΔF508-CFTR-HiBiT(CT) and reduced the ERAD rate by approximately 20% in 293MSR WT cells (Figure 3F). Moreover, the RNF5/185 DKO also decelerated CFTR ERAD and reduced the ERAD rate by approximately 58% (Figure 3F). As expected, the combined HERC3 KD and RNF5/185 DKO resulted in an additive inhibitory effect on CFTR ERAD, reducing its ERAD rate by approximately 75% (Figure 3F). Similar results were obtained in 293MSR, CFBE and BEAS-2B human airway epithelial cells that stably expressed ΔF508-CFTR-Nluc (Taniguchi et al., 2022). HERC3 KD and RNF5/185 double KD (DKD) in these cells additively inhibited CFTR ERAD (Figures 3G, 3H, and 3I). These results collectively indicate that HERC3, in conjunction with RNF5/185, promotes the ERAD of ΔF508-CFTR.

**Figure 3.**
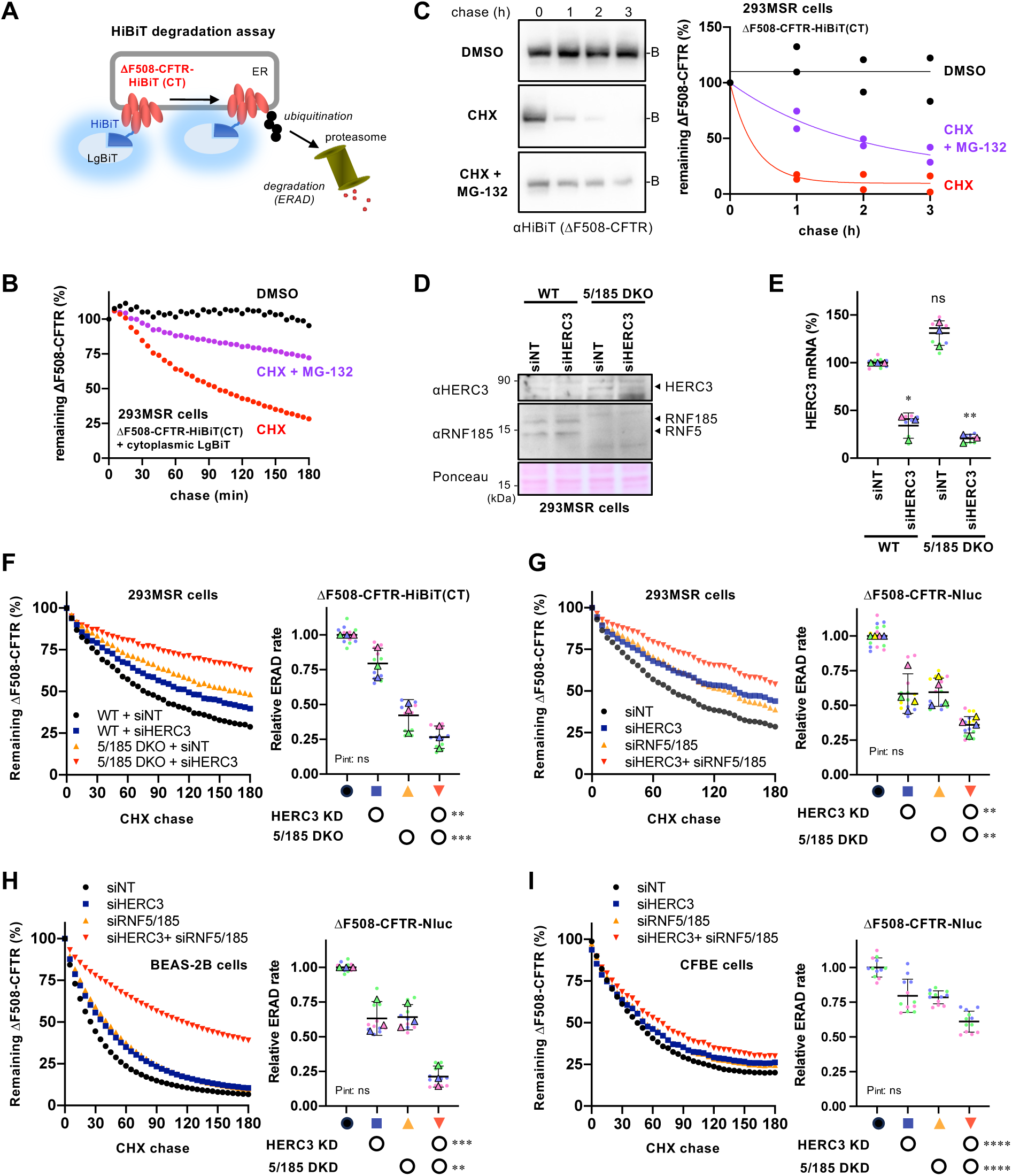
HERC3 and RNF5/185 additively facilitate ΔF508-CFTR ERAD. (A) A schematic diagram of the HiBiT degradation assay, where ΔF508-CFTR-HiBiT(CT) and cytosolic LgBiT were co-expressed. The luminescence signal generated by the interaction of the HiBiT tag and LgBiT was measured in living cells. (B) A typical measurement of ΔF508-CFTR-HiBiT(CT) ERAD in 293MSR cells. The luminescence signal during the CHX chase was measured as the remaining ΔF508-CFTR during the CHX chase, with or without 10 µM MG-132. (C) The metabolic stability of ΔF508-CFTR-HiBiT(CT) was assessed through a CHX chase at 37°C, followed by Western blotting using an anti-HiBiT antibody in 293MSR cells (n=2). The remaining ΔF508-CFTR was expressed as a percentage of time 0, and one-phase exponential decay curves were fitted. (D) Western blotting confirmed the ablation of HERC3, RNF5, and RNF185 in the WT and RNF5/185 DKO 293MSR cells. Ponceau staining was used as a loading control. (E) HERC3 KD in 293MSR WT and RNF5/185 DKO cells was confirmed through quantitative PCR (n=3). (F) Kinetic degradation of ΔF508-CFTR-HiBiT(CT) in 293MSR WT and RNF5/185 KO cells transfected with 50 nM siNT or siHERC3. Luminescence was continuously monitored over 180 minutes in the presence of CHX and plotted normalized to the non-treated cells. The ERAD rate of ΔF508-CFTR-HiBiT(CT) was calculated by fitting the initial degradation portion of each kinetic degradation curve (right, n=3). (G, H, I) Kinetic degradation of ΔF508-CFTR-Nluc(CT) in 293MSR (G, n=4), BEAS-2B (H. n=3), and CFBE (I, n=12) cells transfected with 50 nM siRNA as indicated. The ERAD rate of ΔF508-CFTR-Nluc(CT) was calculated as F. Each biological replicate (n) is color-coded: the averages from 3-4 technical replicates are shown in triangles (E-H). Statistical significance was assessed by one-way RM ANOVA with Dunnett’s multiple comparison tests (E) or two-way RM ANOVA which revealed a significant main effect of HERC3 or RNF5/185 ablation, but no interaction between them (F-I, P_int_> 0.05). Data represent mean ± SD. *p < 0.05, **p < 0.01, ***p < 0.001, ****p < 0.0001, ns, not significant.

### The involvement of HERC3 in the retrotranslocation and ubiquitination of ΔF508-CFTR

To explore how HERC3 facilitates CFTR ERAD, we measured its impact on retrotranslocation which is a crucial step for proteasomal degradation of luminal and membrane proteins (Hampton and Sommer, 2012; Lemberg and Strisovsky, 2021; Wu and Rapoport, 2018). We developed a live cell CFTR retrotranslocation assay using ΔF508-CFTR fused with the HiBiT tag in the extracellular region (ΔF508-CFTR-HiBiT(Ex)) (Taniguchi et al., 2022). The HiBiT tag initially located in the luminal side of the ER is expected to transfer to the cytoplasm where it can associate with co-expressed LgBiT after retrotranslocation (Figure 4A). This results in a reconstituted Nluc luminescence signal that can be measured in real-time in living cells. The luminescence signal was observed exclusively when both ΔF508-CFTR-HiBiT(Ex) and LgBiT were expressed (Figure S2A). Furthermore, in the presence of the proteasome inhibitor MG-132, the luminescence signal exhibited a continuous increase, indicating the cytoplasmic accumulation of ΔF508-CFTR-HiBiT(Ex) (Figures 4B and S2A). Treatment with DBeQ, an inhibitor of p97/VCP, which is crucial for retrotranslocation (Chou et al., 2011), abrogated the luminescence increase induced by MG-132, suggesting that this luminescence signal can be used as an indicator of retrotranslocated CFTR from the ER to the cytoplasm (Figure 4B). Furthermore, we observed an elevation in the luminescent signal during MG-132 treatment, even when CHX was present. This finding suggests that the increased signal is likely attributed to the retrotranslocation of pre-existing ΔF508-CFTR-HiBiT(Ex) located within the ER (Figure 4C). Moreover, we observed a reduced retrotranslocation of ΔF508-CFTR-HiBiT(Ex) upon treatment with the CFTR modulator Trikafta, known for its ability to correct CFTR misfolding and prevent premature degradation (Capurro et al., 2021; Keating et al., 2018) (Figure 4C). The HiBiT retrotranslocation assay showed that consistent with the ERAD inhibitory effect, HERC3 KD tended to slightly reduce the retrotranslocation of ΔF508-CFTR-HiBiT(Ex) in 293MSR WT cells (Figure 4D). As expected, RNF5/185 DKO robustly inhibited the CFTR retrotranslocation, and this effect was additively enhanced by HERC3 KD (Figure 4D). These inhibitory effects on retrotranslocation were highly correlated with the effects on ERAD (Figure S3A).

**Figure 4.**
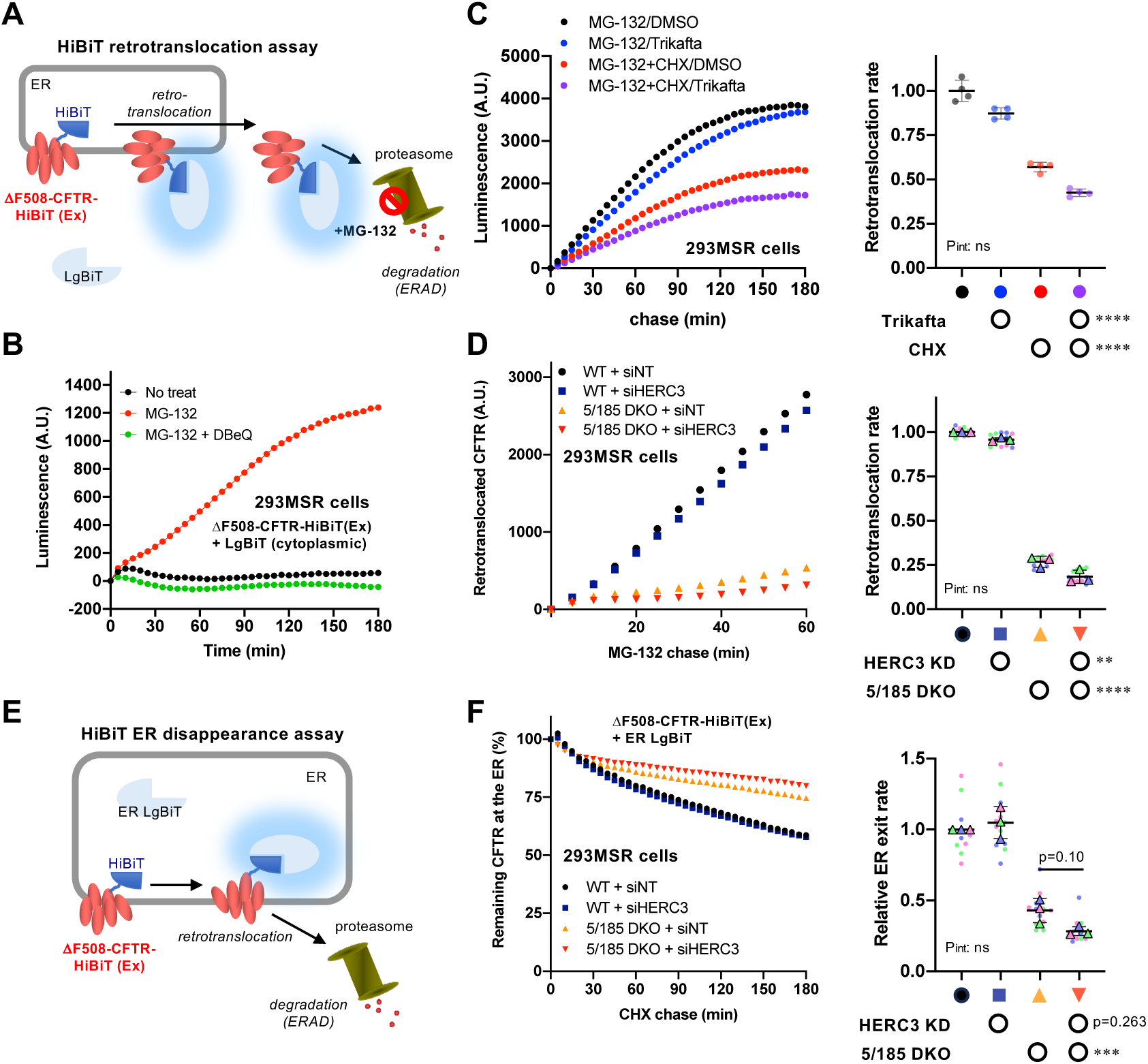
HERC3 and RNF5/185 facilitate ΔF508-CFTR retrotranslocation. (A) A schematic diagram of the HiBiT retrotranslocation assay, where ΔF508-CFTR-HiBiT(Ex) and cytosolic LgBiT were co-expressed. The luminescence signal generated by the interaction of LgBiT and the HiBiT tag exposed in the cytosol after retrotranslocation was measured in living cells during MG-132 treatment. (B) A typical measurement of ΔF508-CFTR-HiBiT(Ex) retrotranslocation in 293MSR cells. The luminescence signal was measured in living cells upon treatment with 10 µM MG-132, with or without 10 µM DBeQ. (C) Kinetic retrotranslocation of ΔF508-CFTR-HiBiT(Ex) in 293MSR cells treated with DMSO (0.3%) or Trikafta (3 µM VX-661, 3 µM VX-445, 1 µM VX-770) for 24 hours at 37°C. Luminescence was continuously monitored in the presence of MG-132 with or without CHX. The signal increased by the MG-132 treatment was plotted as retrotranslocated CFTR. The retrotranslocation rate of ΔF508-CFTR-HiBiT(Ex) was calculated by linear fitting of the signal until 60 min (right, n=4). Two-way RM ANOVA revealed a significant main effect of Trikafta or CHX, but no interaction between them (P_int_>0.05). (D) Kinetic retrotranslocation of ΔF508-CFTR-HiBiT(Ex) in 293MSR WT and RNF5/185 KO cells transfected with 50 nM siNT or siHERC3. Luminescence was continuously monitored over 60 minutes in the presence of MG-132. The signal increased by the MG-132 treatment was plotted as retrotranslocated CFTR. The retrotranslocation rate of ΔF508-CFTR-HiBiT(Ex) was calculated by linear fitting (right, n=3). Two-way RM ANOVA revealed a significant main effect of HERC3 KD or RNF5/185 DKO, but no interaction between them (P_int_>0.05). (E) A schematic diagram of the HiBiT ER disappearance assay, where ΔF508-CFTR-HiBiT(Ex) and ER-luminal LgBiT (ER LgBiT) were co-expressed. The luminescence signal generated by the interaction of LgBiT and the HiBiT tag in the ER was measured in living cells during the CHX chase. (F) Kinetic ER disappearance of ΔF508-CFTR-HiBiT(Ex) in 293MSR WT and RNF5/185 KO cells transfected with 50 nM siNT or siHERC3. Luminescence was continuously monitored over 180 minutes in the presence of CHX and plotted normalized to the non-treated cells as remaining CFTR at the ER (%). The ER disappearance rate of ΔF508-CFTR-HiBiT(Ex) was calculated by fitting the kinetic ER disappearance curve (right, n=3). Two-way RM ANOVA with Holm-Sidak multiple comparison tests revealed a significant main effect of RNF5/185 DKO and no interaction between HERC3 KD and RNF5/185 DKO (P_int_>0.05). Each biological replicate (n) is color-coded: the averages from 3 or 4 technical replicates are shown in triangles (D, F). Data represent mean ± SD. *p < 0.05, **p < 0.01, ***p < 0.001, ****p < 0.0001, ns, not significant.

The ΔF508-CFTR-HiBiT(Ex) retrotranslocation was also measured using co-expressed ER luminal LgBiT (ER LgBiT) which was fused with the ER signal peptides of calnexin (CNX) at the N-terminus, and an ER retention signal (KDEL) at the C-terminus (Figure 4E). Luminescence was measured in real-time after adding CHX, and the decrease in luminescence corresponded to the disappearance of ΔF508-CFTR-HiBiT(Ex) from the ER. This ER disappearance assay couldn’t detect the weak effect of HERC3 KD on the CFTR retrotranslocation in the WT cells (Figure 4F). However, it was able to detect that RNF5/185 DKO reduced the disappearance of ΔF508-CFTR-HiBiT(Ex) from the ER (Figure 4F). Like the results in the HiBiT retrotranslocation assay, HERC3 KD tended to reduce the ER disappearance rate of ΔF508-CFTR in RNF5/185 DKO cells (Figure 4F). The results of both retrotranslocation analyses were highly correlated (Figure S3B). Taken together, our HiBiT assays reveal that HERC3 appears to be involved in ΔF508-CFTR retrotranslocation, albeit slightly, independently of RNF5/185.

Next, we examined whether HERC3 regulates CFTR ubiquitination independently of RNF5/185. Surprisingly, Western blotting with a pan-Ub antibody did not detect a substantial reduction in the total ubiquitination of immature HBH-ΔF508-CFTR upon HERC3 KD in 293MSR WT cells and RNF5/185 DKO cells (Figure 5A). However, RNF5/185 DKO resulted in a significant reduction in the total CFTR ubiquitination (Figure 5A). To obtain more quantitative results, the CFTR ubiquitination level was assessed using Ub ELISA, a sensitive and highly quantitative method (Kamada et al., 2019; Okiyoneda et al., 2018). Ub ELISA showed that HERC3 KD reduced both K48- and K63-linked polyubiquitination of immature HBH-ΔF508-CFTR by approximately 45% in 293MSR WT cells (Figures 5B and 5C). As expected, RNF5/185 DKO reduced both K48- and K63-linked polyubiquitination of immature HBH-ΔF508-CFTR by about 75% (Figures 5B, 5C, and S3C). Interestingly, the effect of HERC3 KD on K48- and K63-linked polyubiquitination was somewhat antagonized in RNF5/185 DKO cells. Nonetheless, in direct comparison, with the use of higher amounts of RNF5/185 DKO cell lysate, the KD of HERC3 still significantly reduced both K48- and K63-linked polyubiquitination of immature ΔF508-CFTR, suggesting that HERC3, to some extent, promotes CFTR ubiquitination independently of RNF5/185 (Figure 5D).

**Figure 5.**
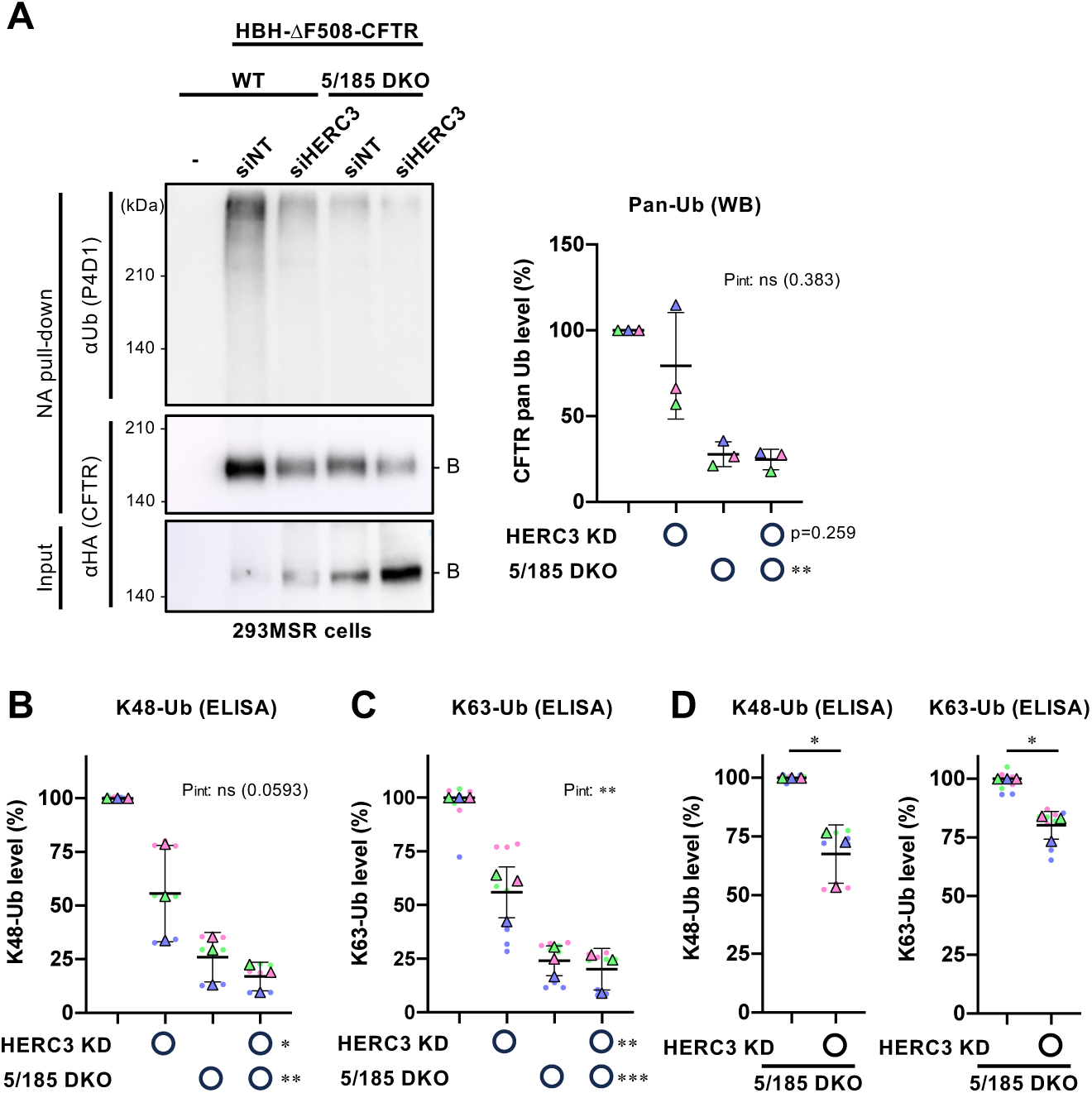
HERC3 and RNF5/185 facilitate ΔF508-CFTR ubiquitination. (A) Ubiquitination levels of HBH-ΔF508-CFTR-3HA in 293MSR WT and RNF5/185 DKO cells were measured by Neutravidin pull-down under denaturing conditions (NA pull-down) and Western blotting. The CFTR ubiquitination level was quantified by densitometry and normalized to CFTR in precipitates (right, n=3). Two-way RM ANOVA revealed a significant main effect of RNF5/185 DKO and no interaction between HERC3 KD and RNF5/185 DKO (P_int_>0.05). (B, C) K48 (B, n=3) and K63-linked poly-ubiquitination (C, n=3) of HBH-ΔF508-CFTR in 293MSR WT and RNF5/185 DKO cells transfected with 50 nM siNT or siHERC3 were quantified by Ub ELISA using Ub linkage-specific antibodies. 10 µM MG-132 was treated for 3 hours at 37˚C. The ubiquitination level was normalized by the CFTR amount quantitated by ELISA using an anti-HA antibody. Two-way RM ANOVA revealed significant main effects of HERC3 KD or RNF5/185 DKO and a significant interaction between them in H, but not in G (P_int_>0.05). (D) The effect of HERC3 KD on K48 and K63-linked poly-ubiquitination of HBH-ΔF508-CFTR in RNF5/185 DKO cells was measured by Ub ELISA using higher amounts of cell lysate. Statistical significance was assessed by paired t-test (n=3). Each biological replicate (n) is color-coded: the averages from 3 or 4 technical replicates are shown in triangles (B, C, D). Data represent mean ± SD. *p < 0.05, **p < 0.01, ***p < 0.001, ****p < 0.0001, ns, not significant.

### HERC3 facilitates the UBQLN2 recruitment to the misfolded CFTR during the ERAD

Given that HERC3 interacts with proteasome shuttling factors UBQLN1 and UBQLN2 (Hochrainer et al., 2008), it is plausible that UBQLN proteins play a role in the HERC3 ERAD branch. Notably, previous studies have indicated that UBQLN1 and UBQLN2 facilitate the ERAD of α1-anti-trypsin null Hong Kong mutant, a misfolded luminal protein, and CD3δ, a membrane-spanning protein (Kim et al., 2008; Lim et al., 2009). These UBQLNs are believed to serve as proteasome shuttling factors that guide ubiquitinated targets to the proteasome (Hjerpe et al., 2016; Itakura et al., 2016). However, their roles in retrotranslocation remain unclear. We particularly focused on UBQLN2 because the overexpression (OE) of UBQLN2, but not UBQLN1, reduced immature ΔF508-CFTR in a dose-dependent manner (Figure 6A). To examine whether HERC3 facilitates the association of ΔF508-CFTR with UBQLN2, pull-down experiments were performed using BHK cells stably expressing HBH-ΔF508-CFTR-3HA. Western blotting demonstrated that HERC3 OE increased the interaction of immature HBH-ΔF508-CFTR-3HA with FLAG-UBQLN2 (Figure 6B). Additionally, an ELISA-based assay was used to quantify the binding of FLAG-UBQLN2 with HBH-ΔF508-CFTR immobilized on NeutrAvidin (NA)-coated plates. In contrast to OE, HERC3 KD modestly reduced the CFTR-UBQLN2 interaction by approximately 35%, while RNF5/185 DKO robustly reduced the interaction by about 72% in 293MSR WT cells (Figure 6C). Like the impact on CFTR ubiquitination, the effect of HERC3 KD on the CFTR-UBQLN2 interaction was antagonized in RNF5/185 DKO cells (Figure 6C). However, when directly compared in RNF5/185 DKO cells transfected with an increased amount of FLAG-UBQLN2, HERC3 KD slightly but significantly reduced the CFTR-UBQLN2 interaction (Figure 6D). A pull-down experiment also confirmed these results, where the association of HBH-ΔF508-CFTR with endogenous UBQLN2 in 293MSR cells was robustly reduced by RNF5/185 DKO and almost undetectable upon HERC3 KD in RNF5/185 DKO cells (Figure 6E). The reduced CFTR-UBQLN2 association upon HERC3 and RNF5/185 ablation was highly correlated with the reduction in CFTR polyubiquitination, suggesting that HERC3 and RNF5/185 promote the CFTR-UBQLN2 interaction mainly by facilitating CFTR ubiquitination (Figures S3H and S3I). Furthermore, these changes in the CFTR-UBQLN2 interaction were also highly correlated with the rates of ΔF508-CFTR ERAD and retrotranslocation, suggesting that the reduced UBQLN2 association could lead to decelerated retrotranslocation and ERAD of misfolded CFTR (Figures S3J and S3K).

**Figure 6.**
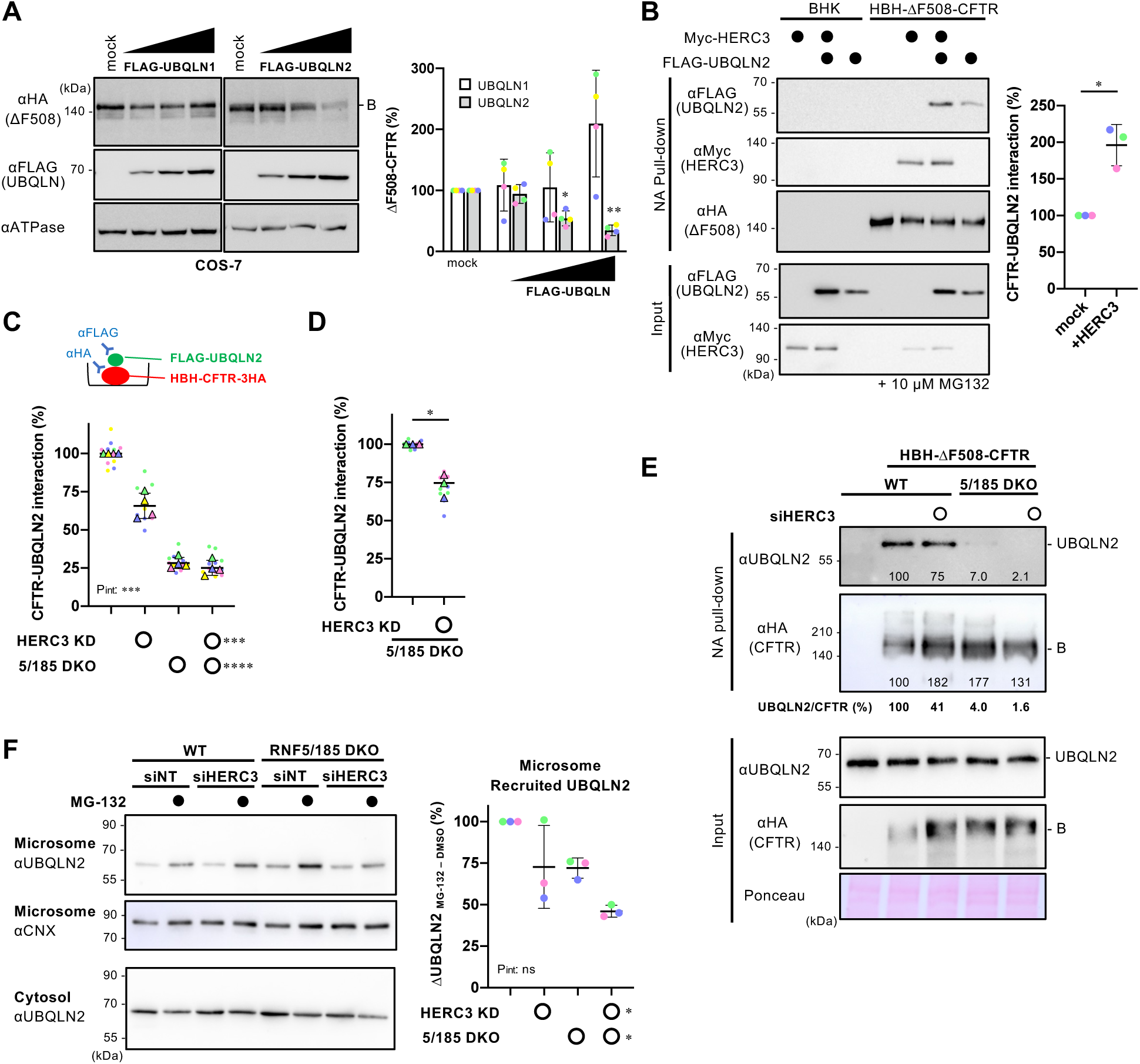
HERC3 facilitates ΔF508-CFTR interaction with UBQLN2. (A) Western blotting showed the steady-state level of ΔF508-CFTR-3HA under OE of FLAG-UBQLN1 or FLAG-UBQLN2 in transiently co-expressed COS-7 cells. The CFTR level was quantified by densitometry (right, n=4). Na^+^/K^+^ ATPase (ATPase) was used as a loading control. B, immature form. (B) The interaction between FLAG-UBQLN2 and HBH-ΔF508-CFTR-3HA in BHK cells transfected with or without Myc-HERC3 was assessed using NA pull-down and Western blotting. The amount of UBQLN2 bound to HBH-ΔF508-CFTR-3HA was quantified by densitometry and normalized to CFTR levels in the precipitates (right, n=3). (C, D) The interaction between FLAG-UBQLN2 and HBH-ΔF508-CFTR-3HA in 293MSR WT and RNF5/185 DKO cells transfected with 50 nM siNT or siHERC3 was measured by ELISA using an anti-FLAG antibody. The level of FLAG-UBQLN2 binding was normalized to the CFTR level, which was measured by ELISA using an anti-HA antibody (C, n=4). Additionally, under conditions of increased FLAG-UBQLN2 expression, the UBQLN2 binding to HBH-ΔF508-CFTR-3HA in RNF5/185 DKO cells was quantified by ELISA (D, n=3). (E) The association of HBH-ΔF508-CFTR with endogenous UBQLN2 in 293MSR WT or RNF5/185 DKO cells transfected with 50 nM siNT or siHERC3 was analyzed by NA pull-down after DSP cross-linking. The quantities of UBQLN2 and ΔF508-CFTR in the precipitates were measured using densitometry and expressed as a percentage of the control. The quantities of CFTR-bound UBQLN2 were normalized to CFTR levels as UBQLN2/CFTR and expressed as a percentage of the control. (F) The level of endogenous UBQLN2 in the microsomes of 293MSR WT and RNF5/185 DKO cells transfected with 50 nM siNT or siHERC3 was measured. Cells were treated with or without 10 µM MG-132 for 3 hours before subcellular fractionation. Microsomes enriched with ER membranes were confirmed using an anti-calnexin (CNX) antibody. The quantities of the ER-recruited UBQLN2 were quantified by subtracting the amount of UBQLN2 before MG-132 treatment from the amount after MG-132 treatment and were expressed as a percentage of the control (n=4, right). Each biological replicate (n) is color-coded: the averages from 3 technical replicates are shown in triangles (D, E). Statistical significance was assessed by one-way RM ANOVA with Dunnett’s multiple comparison tests (A), a paired t-test (B, D), or two-way RM ANOVA (C, F). Data represent mean ± SD. *p < 0.05, ***p < 0.001, ****p < 0.0001, ns, not significant.

During ERAD, cytoplasmic UBQLN proteins are recruited to the ER membrane to aid in the proteasomal degradation of ubiquitinated proteins (Lim et al., 2009). To investigate the role of HERC3 in UBQLN2 recruitment to the ER membrane, we measured the endogenous UBQLN2 abundance in microsomes. Consistent with a previous study (Lim et al., 2009), proteasome inhibitor MG-132 treatment increased the endogenous UBQLN2 in microsomes, indicating the recruitment of UBQLN2 from the cytoplasm to the ER membrane during ERAD (Figure 6F). We quantified the ER-recruited UBQLN2 by measuring the increase in UBQLN2 abundance in the microsome after MG-132 treatment. The results of the quantification showed that the KD of HERC3 and the DKO of RNF5/185 both independently reduced the recruitment of UBQLN2 to the ER membrane. Interestingly, when HERC3 KD and RNF5/185 DKO were combined, there was an additive reduction in the recruitment of UBQLN2 to the ER membrane (Figure 6F). This suggests that HERC3 and RNF5/185 cooperate to promote UBQLN2 recruitment during ERAD.

### UBQLN proteins facilitate the retro-translocation of misfolded CFTR

Next, we determined whether UBQLN2 plays a role in facilitating the retrotranslocation and ERAD of ΔF508-CFTR, as its function in retrotranslocation has not been clearly understood despite its role as proteasome shuttling factors (Hochrainer et al., 2008). Initially, a single KD of UBQLN1, UBQLN2, or UBQLN4 did not significantly reduce ΔF508-CFTR ERAD, possibly due to functional redundancy among UBQLN proteins, which are widely expressed in all tissues (Marin, 2014)(Figure S4A). Therefore, a triple KD of UBQLN1/2/4 was performed and confirmed by Western blotting (Figure 7A). The triple KD of UBQLNs reduced ΔF508-CFTR ERAD by approximately 23% and retrotranslocation by approximately 42% in 293MSR cells (Figures 7B and 7C).

**Figure 7.**
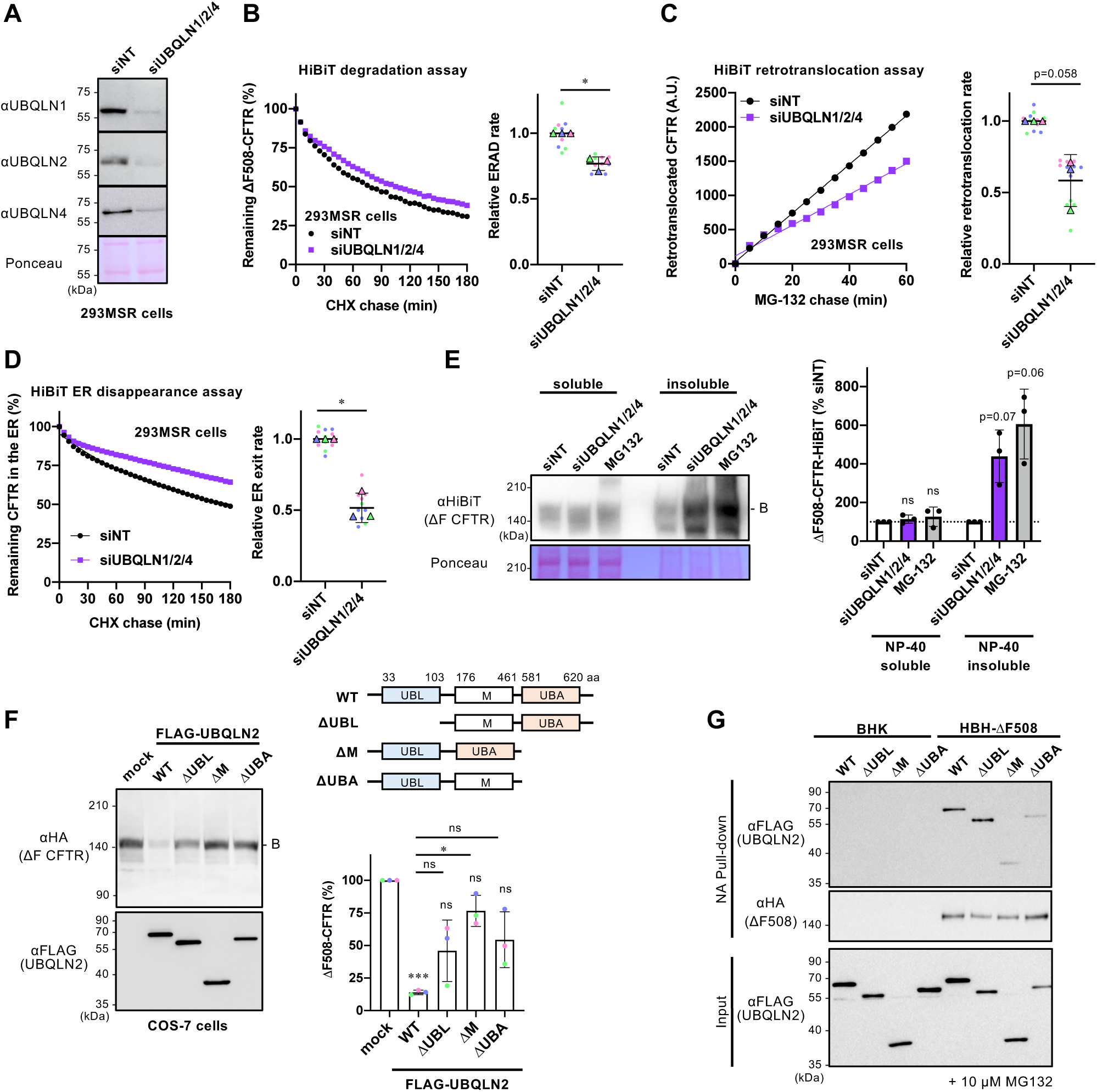
UBQLN proteins facilitate ΔF508-CFTR retrotranslocation and ERAD. (A) Western blotting confirmed the triple KD of UBQLN1, 2, and 4 in 293MSR cells transfected with 50 nM siRNA as indicated. Ponceau staining was used as a loading control. (B) Kinetic degradation of ΔF508-CFTR-HiBiT(CT) in 293MSR WT cells transfected with 50 nM siNT or siUBQLN1/2/4. The ERAD rate was calculated by fitting the initial degradation portion of each kinetic degradation curve (right, n=3). (C) Kinetic retrotranslocation of ΔF508-CFTR-HiBiT(Ex) in 293MSR cells upon UBQLN triple KD. The retrotranslocation was calculated by linear fitting (right, n=3). (D) Kinetic ER disappearance of ΔF508-CFTR-HiBiT(Ex) in 293MSR cells upon UBQLN triple KD. The ER disappearance rate was calculated by fitting the kinetic ER disappearance curve (right, n=3). (E) The detergent NP-40 solubility of ΔF508-CFTR-HiBiT(CT) in 293MSR cells was assessed following UBQLN1/2/4 triple KD or MG-132 treatment (10 µM, 3 hours) using Western blotting with an anti-HiBiT antibody (n=3). The soluble (100 µg) and insoluble (40 µg) fractions were analyzed. (F) The effects of overexpressed FLAG-UBQLN2 variants on the steady-state level of ΔF508-CFTR-3HA were analyzed by Western blotting in co-transfected COS-7 cells. The immature ΔF508-CFTR (B band) was quantified by densitometry (right, n=3). A schematic diagram of the UBQLN2 domain composition with the residue numbers at the domain boundaries. UBQLN2 mutants used in this study are also shown. (G) The interaction of FLAG-UBQLN2 variants with HBH-ΔF508-CFTR-3HA in BHK cells was analyzed by NA pull-down and Western blotting. Cells were treated with 10 µM MG-132 for 3 hours before cell lysis. Statistical significance was assessed by a paired t-test (B, C, D), or one-way RM ANOVA with Dunnett’s multiple comparison tests (E, F). Each biological replicate (n) is color-coded: the averages from 4 technical replicates are shown in triangles (B, C, D). Data represent mean ± SD. *p < 0.05, ns, not significant.

Likewise, we observed reduced retrotranslocation upon UBQLN triple KD even in experiments using MG-132 and CHX chase, indicating that UBQLNs facilitate retrotranslocation of pre-existing ΔF508-CFTR-HiBiT within the ER (Figure S4B). Furthermore, the triple KD of UBQLNs resulted in a reduction of ΔF508-CFTR disappearance from the ER lumen (Figure 7D). These findings suggest that UBQLNs not only promote ERAD but also facilitate the retrotranslocation of misfolded CFTR from the ER to the cytoplasm. The detergent solubility analyses revealed that UBQLN triple KD showed a tendency to increase the insolubility of ΔF508-CFTR-HiBiT(CT) in 293MSR cells, a phenotype similar to that observed with MG-132 treatment (Figure 7E). This suggests that UBQLNs may play a crucial role in maintaining the solubility of ΔF508-CFTR, thereby facilitating retrotranslocation and ERAD. Alternatively, the reduced retrotranslocation and ERAD observed upon UBQLNs KD could lead to increased CFTR aggregation.

To gain further insight into the mechanism of UBQLNs’ action in CFTR ERAD, we tested the effects of UBQLN2 mutants with the Ub-like domain (UBL), central M domain (M), or Ub-associated domain (UBA) deleted (Figure 7F). The UBL and UBA domains are known to be involved in proteasome binding (Chen et al., 2016; Ko et al., 2004) and poly-Ub chain binding (Zhang et al., 2008), respectively, while the central M domain containing stress-inducible 1 (STI1) domain is believed to bind to exposed transmembrane segments in the cytosol to prevent aggregation (Itakura et al., 2016). Western blotting showed that FLAG-UBQLN2 OE reduced immature ΔF508-CFTR, likely due to facilitating ERAD (Figure 7F). In contrast, the deletion of UBL or UBA slightly reduced the effect of UBQLN2, suggesting that UBQLN2 likely recognizes poly-Ub chains on ΔF508-CFTR through its UBA domain and transfers to the proteasome via its UBL domain during ERAD (Figure 7F). Interestingly, the deletion of the central M domain reduced the UBQLN2 effect, indicating the possibility that the M domain might be crucial in shielding the exposed CFTR transmembrane segments in the cytosol to promote ΔF508-CFTR retrotranslocation and ERAD (Figure 7F). To investigate the interaction between UBQLN2 and ΔF508-CFTR during ERAD, we assessed the CFTR-UBQLN2 association in BHK cells following treatment with the proteasome inhibitor MG-132. Pull-down experiments revealed that the deletion of the M domain or UBA domain reduced the association of HBH-ΔF508-CFTR with FLAG-UBQLN2, suggesting the possibility that during ERAD, the M domain and UBA domain might interact with the exposed transmembrane segments and poly-Ub chains in ΔF508-CFTR, respectively (Figure 7G). On the other hand, the UBL domain appeared to be dispensable for the CFTR interaction (Figure 7G) and was likely involved in CFTR ERAD by binding to proteasomes.

### HERC3 selectively facilitates ERAD of misfolded CFTR

To investigate the substrate selectivity of HERC3, we tested the effect of HERC3 KD on several ERAD models including TCRα (an ERAD-Lm substrate (Horimoto et al., 2013)), Insig-1 (an ERAD-M substrate (Lee et al., 2006; Leto et al., 2019), and D18G-TTR (an ERAD-L substrate (Sato et al., 2012; Sato et al., 2007) (Figure 8A). Kinetic ERAD of TCRα-HiBiT and Insig-1-HiBiT were successfully measured in 293MSR cells co-transfected with cytoplasmic LgBiT as a proteasome inhibitor blocked their degradation (Figures S4C and S4C). The HiBiT degradation assay showed that HERC3 KD and/or RNF5/185 DKO did not lead to a reduction in the ERAD rates of TCRα and Insig-1, indicating that neither HERC3 nor RNF5/185 is involved in their ERAD processes (Figures 8B and 8C). Similarly, the KD of HERC3 and/or the DKO of RNF5/185 did not result in a decrease in D18G-TTR ERAD (Figure 8D). In line with the KD phenotypes, the OE of HERC3 had no impact on the steady-state levels of TCRα-HA and FLAG-D18G-TTR in COS-7 cells. However, HERC3 OE resulted in a dose-dependent reduction in the levels of ΔF508-CFTR (Figure 8E). Notably, unlike WT-CFTR, HERC3 OE did not affect the levels of normally folded WT-CFTR, suggesting that HERC3 may specifically facilitate the degradation of misfolded or structurally abnormal CFTR (Figure 8E). We additionally investigated the influence of HERC3 KD on the ERAD of an ABC transporter ABCB1 (MDR1/P-glycoprotein) with a ΔY490 mutation (ΔY490-ABCB1), which is analogous to the ΔF508 mutation in CFTR (Hoof et al., 1994). Since both ΔF508-CFTR and ΔY490-ABCB1 exhibit folding defects in the cytosolic NBD1 region, the ERAD-C pathway might be involved in their degradation, based on previous studies in yeast (Gnann et al., 2004; Nakatsukasa et al., 2008). The HiBiT degradation assay showed that like ΔF508-CFTR, the ERAD of ΔY490-ABCB1 was decelerated in RNF5/185 DKO cells compared to the WT cells (Figure 8F). However, HERC3 KD did not result in a deceleration of ΔY490-ABCB1 ERAD in both WT and RNF5/185 DKO cells (Figure 8F). These findings indicate that RNF5/185 is responsible for the degradation of both ΔF508-CFTR and ΔY490-ABCB1. In contrast, HERC3 may selectively recognize specific molecular determinants that are present only in misfolded CFTR, but not in the ABCB1 mutant.

**Figure 8.**
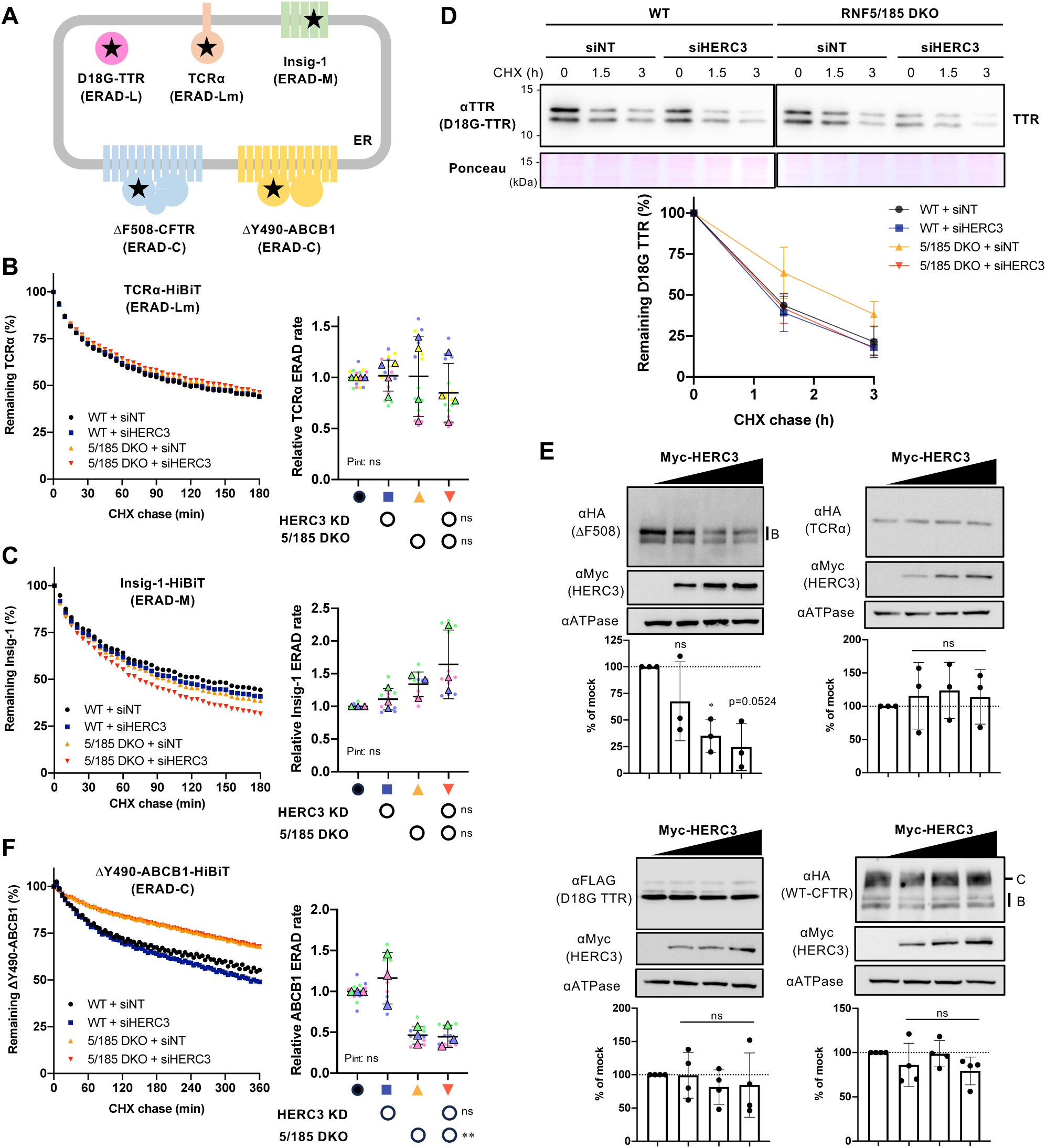
The substrate selectivity of HERC3 in ERAD. (A) A schematic diagram of the ERAD substrate models used in this study. The misfolded region is indicated by a star. The HiBiT tag was fused in the cytoplasmic region except for D18G-TTR. (B, C) The HiBiT degradation assay measured the ERAD of TCRα-HiBiT (B, n=4) and Insig-1-HiBiT (C, n=3) in 293MSR WT and RNF5/185 KO cells transfected with 50 nM siNT or siHERC3, as indicated. (D) The metabolic stability of D18G-TTR was measured by CHX chase at 37°C and Western blotting with an anti-TTR antibody in 293MSR WT and RNF5/185 KO cells transfected with 50 nM siNT or siHERC3 as indicated. The remaining TTR was quantified by densitometry and expressed as a percentage of the initial amount (right, n=3). (E) Western blotting analyzed the effects of Myc-HERC3 OE on co-transfected ΔF508-CFTR-3HA, TCRα-HA, D18G-TTR-FLAG, or WT-CFTR-3HA in COS-7 cells. The immature ΔF508-CFTR (B band), TCRα, D18G-TTR, and total WT-CFTR (B and C bands) were quantified by densitometry (n=3). (F) The HiBiT degradation assay measured the ERAD of ΔY490-ABCB1-HiBiT (E, n=3) in 293MSR WT and RNF5/185 KO cells as B and C. Each biological replicate (n) is color-coded: the averages from 4 technical replicates are shown in triangles (B, C, F). Statistical significance was assessed by a one-way RM ANOVA with Dunnett’s multiple comparison tests (E) or two-way RM ANOVA revealed no significant main effect of HERC3 KD or RNF5/185 DKO, and no interaction between them (P_int_>0.05), except for a significant main effect of RNF5/185 DKO in F. Data represent mean ± SD. *p < 0.05, **p < 0.01, ns, not significant.

To investigate the molecular determinants crucial for HERC3 interaction, we generated CFTR fragments fused with the HiBiT tag at the C-terminus, located in the cytoplasm (Figure 9A). By utilizing the HiBiT degradation assay, we measured the contributions of HERC3 and RNF5/185 to the ERAD of these CFTR fragments. Consistent with previous findings (Du and Lukacs, 2009), MSD1 (M1), NBD1 with the ΔF508 mutation (ΔF508-NBD1), MSD1-NBD1 with ΔF508 mutation (M1-N1(ΔF)), and MSD2 (M2) fragments were rapidly eliminated, indicating that individually expressed CFTR domains are recognized as nonnative polypeptides by the ERQC mechanism (Figures 9B, 9C, 9D, and S5A). Like the full-length ΔF508-CFTR, HERC3 KD and RNF5/185 DKO additively reduced the ERAD rates of M1, M1-N1(ΔF), and M2 fragments in 293MSR cells (Figures 9B, 9C, and 9D). In contrast, the KD of HERC3 and/or RNF5/185 DKO did not have any effect on the ERAD of the cytoplasmic ΔF508-NBD1 (Figures S5A). These findings suggest that HERC3 and RNF5/185 play a role in identifying structural abnormalities in the MSDs of CFTR, thereby aiding in the ERAD of misfolded CFTR. This interpretation gains support from the results showing that HERC3 KD and RNF5/185 DKO had an additional impact on reducing the ERAD rate of N1303K-CFTR (Figure S5B). N1303K-CFTR is known for its NBD2 mutation, which leads to the unfolding of MSD1 and MSD2, as evidenced by limited protease susceptibility (Du and Lukacs, 2009). Furthermore, correlation analyses revealed that the impact of HERC3 and/or RNF5/185 ablation on ERAD was almost equivalent between ΔF508-CFTR and M1 (Figure S5C, slope 0.81) or M1-N1 (Figure S5D, slope 1.06), indicating that HERC3 and RNF5/185 might primarily sense conformational defects in the N-terminal region of CFTR, which is crucial for ΔF508-CFTR ERAD. In contrast, the effect on the ERAD of M2 was weaker compared to that of ΔF508-CFTR (Figure S5E, slope 0.532). These results suggest that in addition to RNF5/185 and HERC3, other E3 ligases, such as CHIP, may also participate in the ERQC checkpoints of MSD2 at the late stages of CFTR biogenesis as proposed previously (Younger et al., 2006).

**Figure 9.**
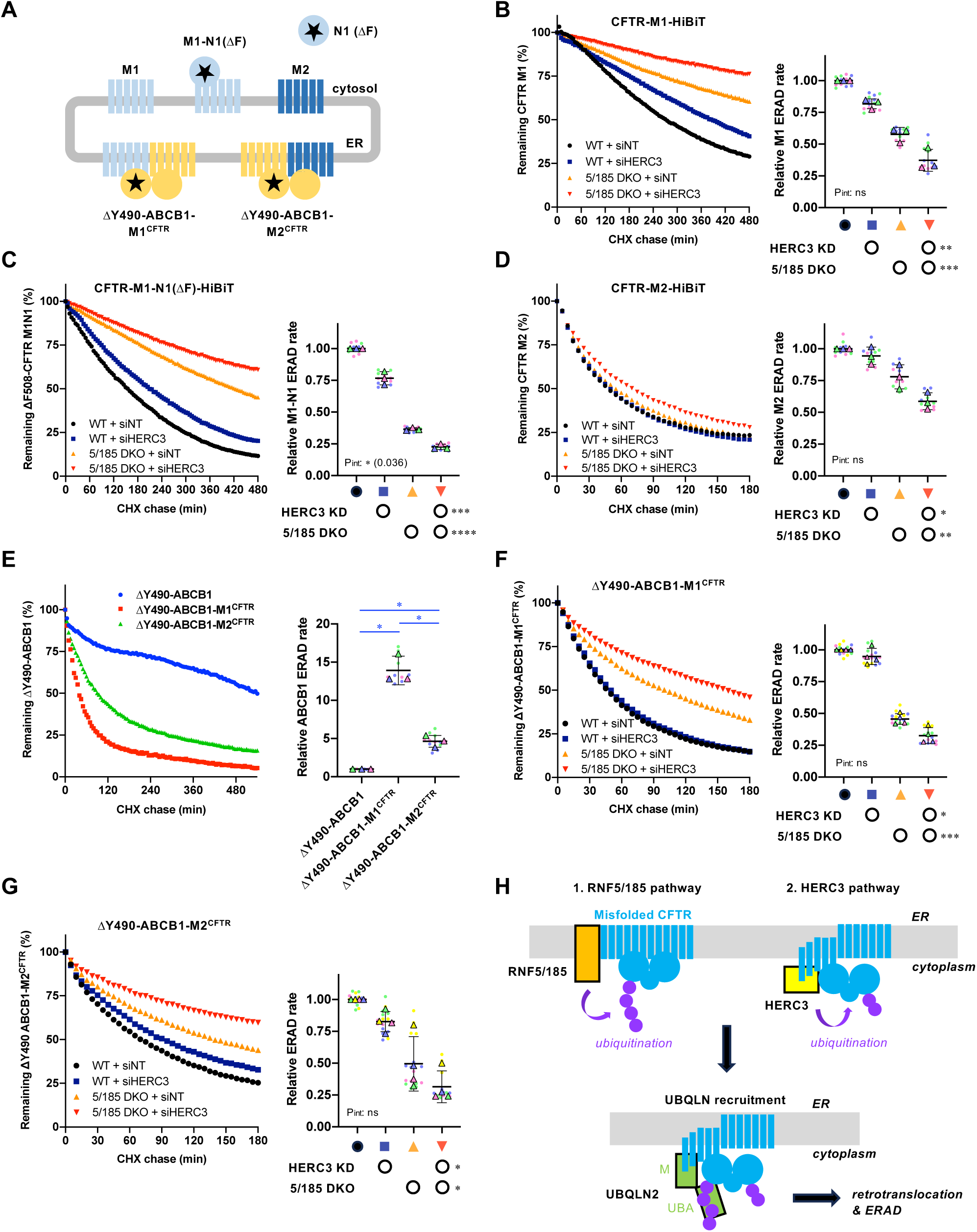
HERC3 selectively facilitates ERAD by interacting with the CFTR-MSDs. (A) A schematic diagram of the CFTR fragment models used in this study. The misfolded region is indicated by a star. M1; MSD1, M1-N1(ΔF); MSD1 and NBD1 with ΔF508 mutation, M2; MSD2, N1(ΔF); NBD1 with ΔF508 mutation. ΔY490-ABCB1-MSD1^CFTR^ and ΔY490-ABCB1-MSD2^CFTR^ are the chimeras in which the MSD1 and MSD2 of ABCB1 were replaced with respective MSDs of CFTR. The HiBiT tag was fused in the C-terminal region located in the cytoplasm. (B, C, D) The HiBiT degradation assay measured the ERAD of M1-HiBiT (B, n=3), M1-N1(ΔF)-HiBiT (C, n=3), and M2-HiBiT (D, n=3) in 293MSR WT and RNF5/185 KO cells transfected with 50 nM siNT or siHERC3, as indicated. (E) The HiBiT degradation assay measured the ERAD of ΔY490-ABCB1, ΔY490-ABCB1-M1^CFTR^, and ΔY490-ABCB1-M2^CFTR^ in 293MSR WT cells (n=3). (F, G) The HiBiT degradation assay measured the ERAD of ΔY490-ABCB1-M1^CFTR^ (F, n=4) and ΔY490-ABCB1-M2^CFTR^ (G, n=4) in 293MSR WT and RNF5/185 KO cells transfected with 50 nM siNT or siHERC3, as indicated. Statistical significance was assessed using one-way RM ANOVA with Dunnett’s multiple comparison tests (E) or two-way RM ANOVA (B, C, D, F, G) which revealed a significant main effect of HERC3 KD or RNF5/185 DKO, but no significant interaction between them (Pint > 0.05), except for C. Each biological replicate (n) is color-coded, and the averages from four technical replicates are represented by triangles. Data represent mean ± SD. *p < 0.05, **p < 0.01, ***p < 0.001, ****p < 0.0001, ns, not significant. (H) The proposed model illustrates the function of HERC3 in the CFTR ERQC. In addition to RNF5/185, HERC3 senses the MSDs of misfolded CFTR. The cytosolic HERC3 interacts with transmembrane domains exposed to the cytoplasm. The RNF5/185- and HERC3-mediated polyubiquitination facilitates UBQLN2 recruitment, thereby promoting CFTR retrotranslocation and ERAD.

To determine whether the CFTR MSDs play a crucial role in HERC3- and RNF5/185-mediated ERAD, we conducted domain-swapped experiments. We created ΔY490-ABCB1-M1^CFTR^ and ΔY490-ABCB1-M2^CFTR^, where the MSD1 or MSD2 of ABCB1 was replaced with the respective CFTR’s MSD (Figure 9A). Using the HiBiT degradation assay, we observed a significantly faster ERAD rate in the ΔY490-ABCB1-M1^CFTR^ and ΔY490-ABCB1-M2^CFTR^ chimeras compared to ΔY490-ABCB1, indicating the presence of potential degrons in the CFTR-MSDs that promote ERAD (Figures 9E). Notably, HERC3 KD and RNF5/185 DKO additively attenuated the ERAD of both ΔY490-ABCB1-M1^CFTR^ and ΔY490-ABCB1-M2^CFTR^ chimeras, in contrast to the observed effect on ΔY490-ABCB1 (Figures 9F and 9G). These findings strongly suggest that HERC3 selectively identifies specific features within CFTR’s MSDs, thereby facilitating the ERAD of specific membrane proteins.

## Discussion

In this study, we have identified HERC3 as an auxiliary E3 ligase that operates alongside RNF5/185 in the ERQC of membrane proteins. Additionally, we have developed HiBiT-based assays to monitor the kinetic ERAD and retrotranslocation of various ERAD substrates. While previous studies used split Venus (Grotzke et al., 2013) or GFP (Zhong et al., 2015) for similar assays, our method offers advantages. We use luminescence, which is more sensitive and suitable for screening chemicals that might have autofluorescence. Furthermore, our assay enables real-time analysis of live cells in a 96-well plate using a plate reader, providing high temporal resolution and the capability to analyze multiple samples, making it a superior choice. By utilizing the HiBiT and Nluc technologies for quantitative retrotranslocation and ERAD measurements, we have discovered that HERC3, in parallel with the RNF5/185 pathway, accelerates the retrotranslocation and ERAD of misfolded CFTR by promoting polyubiquitination. Both HERC3 and RNF5/185 appear to collaborate in recruiting UBQLNs, which facilitates the retrotranslocation and proteasomal delivery of ubiquitinated CFTR. While RNF5/185 contributes to both retrotranslocation and the proteasomal degradation of ΔF508-CFTR, the role of HERC3 in retrotranslocation is relatively minor. HERC3 likely plays a more significant role in facilitating the proteasomal degradation of CFTR following retrotranslocation. Considering that HERC3 is primarily localized in the cytoplasm (Cruz et al., 2001), it is reasonable that HERC3 primarily acts on the misfolded CFTR dislocated to the cytoplasm. Moreover, this presumed function is in line with previous findings that HERC3 promotes the proteasomal degradation of cytosolic proteins such as RPL23A (Zhang et al., 2022b), EIF5A (Zhang et al., 2022a), and SMAD7 (Li et al., 2019). On the other hand, HERC3 appears to be at least partially involved in retrotranslocation, and its contribution may be particularly important when the function of RNF5/185 is insufficient. We speculate that in the absence of RNF5/185, other ER-based Ub ligases might contribute to facilitating CFTR retrotranslocation, especially preceding the HERC3 ERAD pathway. Further research in the future will be necessary to confirm this hypothesis.

Our findings indicate that HERC3 plays a role in facilitating both K48- and K63-linked polyubiquitination of ΔF508-CFTR (Figure S3C). The significant correlation between the reduced polyubiquitination and ERAD upon HERC3 KD suggests that HERC3 promotes the CFTR ERAD by enhancing polyubiquitination (Figures S3D and S3E). In contrast, HERC3 KD led to a ∼45% decrease in the CFTR polyubiquitination, while only marginally affecting the CFTR retrotranslocation (Figures S3F and S3G). This may also support our model that HERC3-mediated ubiquitination primarily influences the downstream ERAD process rather than retrotranslocation. Previous studies have suggested that polyubiquitin chains on ERAD substrates undergo trimming or complete removal by p97/Cdc48-bound deubiquitinases during their extraction from the ER membrane (Ernst et al., 2009; Stein et al., 2014). Additionally, the extracted misfolded membrane proteins promptly interact with the cytosolic E3 ligase RNF126 which catalyzes the reubiquitination of BAG6-captured membrane proteins, facilitating their efficient proteasomal degradation (Hu et al., 2020). Like RNF126, it is plausible that HERC3 also participates in the reubiquitination process of extracted misfolded membrane proteins in the cytoplasm. However, HERC3 seems to at least play a role in the regulation of CFTR ubiquitination at the ER before retrotranslocation, as evidenced by the fact that its KD led to an increase in the abundance of the foldable ΔF508-CFTR biogenic intermediate which can reach the PM, especially at low temperatures or in the presence of a CFTR corrector. In contrast to HERC3, RNF5/185 appears to primarily facilitate retrotranslocation by promoting K48- and K63-linked polyubiquitination (Figures S3C-G). Surprisingly, while HERC3 KD showed an additive effect on reducing ERAD and retrotranslocation when combined with RNF5/185 DKO, the effect of HERC3 KD on CFTR K48- and K63-linked polyubiquitination was counteracted by RNF5/185 DKO. Thus, it appears that the HERC3-mediated CFTR ubiquitination is at least partially dependent on RNF5/185. Similar to Gp78 (Morito et al., 2008), HERC3 may function as an E4 enzyme that elongates the Ub chains initiated by RNF5/185, as it directly binds to Ub (Cruz et al., 2001). Alternatively, HERC3 may modify the Ub chains initiated by RNF5/185 by adding non-canonical Ub chains, such as K27 chains, which promote the p97-proteasome pathway (Shearer et al., 2022), as HERC3 has been shown to promote K27-linked polyubiquitination (Zhang et al., 2022a).

Based on our findings, it appears that HERC3 and RNF5/185 cooperatively facilitate the recruitment of UBQLN2 to the misfolded CFTR at the ER membrane, likely by enhancing CFTR polyubiquitination. In addition to their role as proteasome shuttling factors (Hjerpe et al., 2016; Itakura et al., 2016), our HiBiT-based retrotranslocation assays have revealed a novel function of UBQLNs in promoting retrotranslocation. This novel function of UBQLNs in retrotranslocation aligns with previous findings that the depletion of UBQLNs leads to ER stress, which arises from the accumulation of abnormal proteins in the ER (Lim et al., 2009). While UBQLNs appear to bind ubiquitinated CFTR through their UBA domain, the central M domain of UBQLNs may shield the exposed CFTR transmembrane segments of the dislocation intermediate as proposed (Itakura et al., 2016). Structural models also suggest that the M domain forms a hydrophobic groove that could potentially recognize the transmembrane segments (Fry et al., 2021). Considering that the hydrophobicity of transmembrane helices presents an energetic barrier during the retrotranslocation of integral membrane ERAD substrates (Guerriero et al., 2017), UBQLNs may serve to protect the exposed transmembrane segments of the dislocation intermediate from undesired interactions or aggregation on the surface of the ER membrane, thereby facilitating the retrotranslocation. This model is supported by our observation that UBQLNs triple KD increased the insolubility of ΔF508-CFTR. UBQLNs may also facilitate retrotranslocation through a polyubiquitin-mediated ratcheting mechanism, as proposed previously (Baldridge and Rapoport, 2016). Recently, a transmembrane ER protein TMUB1, which is part of the RNF185 complex (van de Weijer et al., 2020), has been identified as an ER-resident escortase that facilitates the p97-mediated extraction of membrane proteins (Wang et al., 2022). TMUB1 is believed to selectively escort hydrophobic transmembrane segments, but not those with relatively low hydrophobicity (Wang et al., 2022). This characteristic of TUMB1 contrasts with that of UBQLNs, which favors slightly lower hydrophobic transmembrane segments (Itakura et al., 2016). While it remains uncertain whether TMUB1 is implicated in CFTR ERAD, future studies examining the potential roles of the RNF185-TMUB1 axis and the HERC3-UBQLNs axis will be crucial in elucidating whether these pathways have distinct or overlapping functions in retrotranslocation and ERAD. Such investigations will contribute to a comprehensive understanding of the ERQC mechanisms involved in membrane proteins.

Our kinetic ERAD assays, utilizing a variety of substrates, have revealed the impressive substrate specificity of HERC3. It appears that neither the HERC3 nor RNF5/185 ERAD pathways are involved in the degradation of TCRα, Insig-1, and D18G-TTR. These substrates are primarily targeted by the Hrd1 (Kikkert et al., 2004; Sato et al., 2012) and/or Gp78 ERAD branches (Chen et al., 2006; Song et al., 2005). Unlike a cytosolic E3 ligase CHIP involved in the ubiquitination of conformationally defective cytoplasmic NBD1 (Meacham et al., 2001; Rabeh et al., 2012; Younger et al., 2004), HERC3 does not play a crucial role in the QC checkpoint of the CFTR’s NBD1. Consistent with previous studies (van de Weijer et al., 2020; Zhong et al., 2009), the RNF5/185 ERAD branch appears to specifically eliminate certain types of membrane proteins, such as ΔF508-CFTR and ΔY490-ABCB1. Both CFTR and ABCB1 belong to the ABC transporter family, and conserved residues in these ABC transporters are primarily located in the cytosolic region, while sequences in the transmembrane region show high variability (Liu et al., 2017). Like the ΔF508 mutation in CFTR, the ΔY490 mutation causes misfolding of ABCB1, disrupting the packing of the transmembrane segments (Loo et al., 2002). Therefore, RNF5/185 appears to play a role primarily in the ERQC checkpoint of the MSDs in mutants of both ABCB1 and CFTR, as proposed previously (Younger et al., 2006). Based on our results using the ABCB1-CFTR chimera, it seems that the CFTR’s MSD1 and MSD2 may contain degrons that facilitate the ERAD of polytopic membrane protein. This finding aligns with recent research showing that a type I CFTR corrector, tezacaftor (VX-661), and a type III corrector, elexacaftor (VX-445), bind to MSD1 and MSD2, respectively, stabilizing these domains and reducing the susceptibility of mutant CFTR to degradation by the ERQC machinery (Fiedorczuk and Chen, 2022b). Since introducing the CFTR’s MSD1 or MSD2 was sufficient to induce the HERC3-dependent ERAD of ΔY490-ABCB1, HERC3 likely identifies specific features of conformationally defective CFTR’s MSDs through its N-terminal RLD, known to be involved in substrate recognition in other HERC families (Garcia-Gonzalo and Rosa, 2005; Hadjebi et al., 2008; Yagita et al., 2023). In contrast to ABCB1 transmembrane segments which lack charged residues (Wang et al., 2007), the insertion and orientation of CFTR transmembrane segments could be more error-prone due to the presence of a substantial number of charged residues in these segments (Sadlish and Skach, 2004). Specifically, studies have demonstrated that transmembrane segment 6 of CFTR, which contains three positively charged residues, is highly unstable in the lipid bilayer (Tector and Hartl, 1999). Consequently, the MSDs of CFTR are predicted to be unstable in the membrane (Fiedorczuk and Chen, 2022a). We speculate that in addition to the ER-embedded RNF5/185, the cytosolic E3 ligase HERC3 provides an ERAD branch specifically recognizing the exposed degron in the less hydrophobic MSDs of CFTR on the cytoplasmic surface of the ER membrane (Figure 9H).

## Acknowledgments

We thank R. Kopito and Y. Ye (TCRα-HA), M. Gottesman (Addgene #10957 MDRwt /ABCB1), P. Howley (Addgene #8661 FLAG-hPLIC-2/UBQLN2, Addgene #8663 FLAG-hPLIC-1/UBQLN1), H. Kai (D18G-TTR), D. Root (Addgene #41394 pLIX402, #25890 pLX304), F. Zhang (Addgene #62988 pSpCas9(BB)-2A-Puro (PX459) V2.0) for providing plasmids, D. Gruenert (University of California, San Francisco) for the CFBE41o-cells, S. Kato for technical support.

## Author Contributions

Yuka Kamada: Methodology, Investigation, Writing draft, Yuko Ohnishi, Chikako Nakashima, Aika Fujii, Mana Terakawa, Ikuto Hamano, Uta Nakayamadam, Saori Katoh, Noriaki Hirata, Hazuki Tateishi, Ryosuke Fukuda: Methodology, Investigation, Hirotaka Takahashi: Resources, Gergely L. Lukacs: Conceptualization, Resources, Tsukasa Okiyoneda: Conceptualization, Formal analysis, Writing, Visualization, Supervision, Project administration, Funding acquisition.

## Lead contact

Further information and requests for resources and reagents may be directed to and will be fulfilled by Tsukasa Okiyoneda.

## Materials availability

Plasmids generated in this study are available by request from the lead contact with a Material Transfer Agreement with Kwansei Gakuin University.

## Data Availability

The raw data required to reproduce these findings are available from the corresponding authors upon reasonable request.

## Materials and Methods

### KEY RESOURCE TABLE

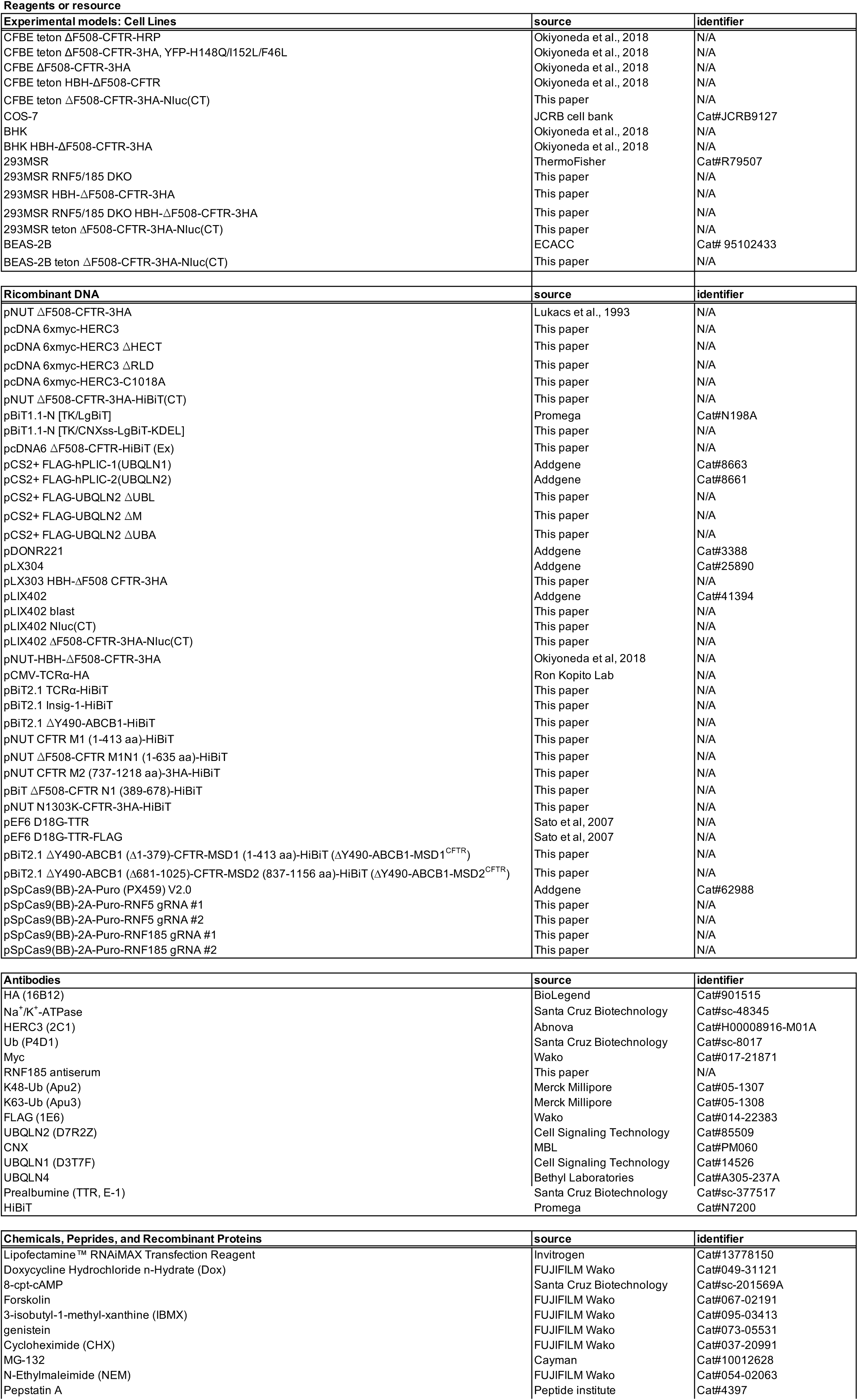
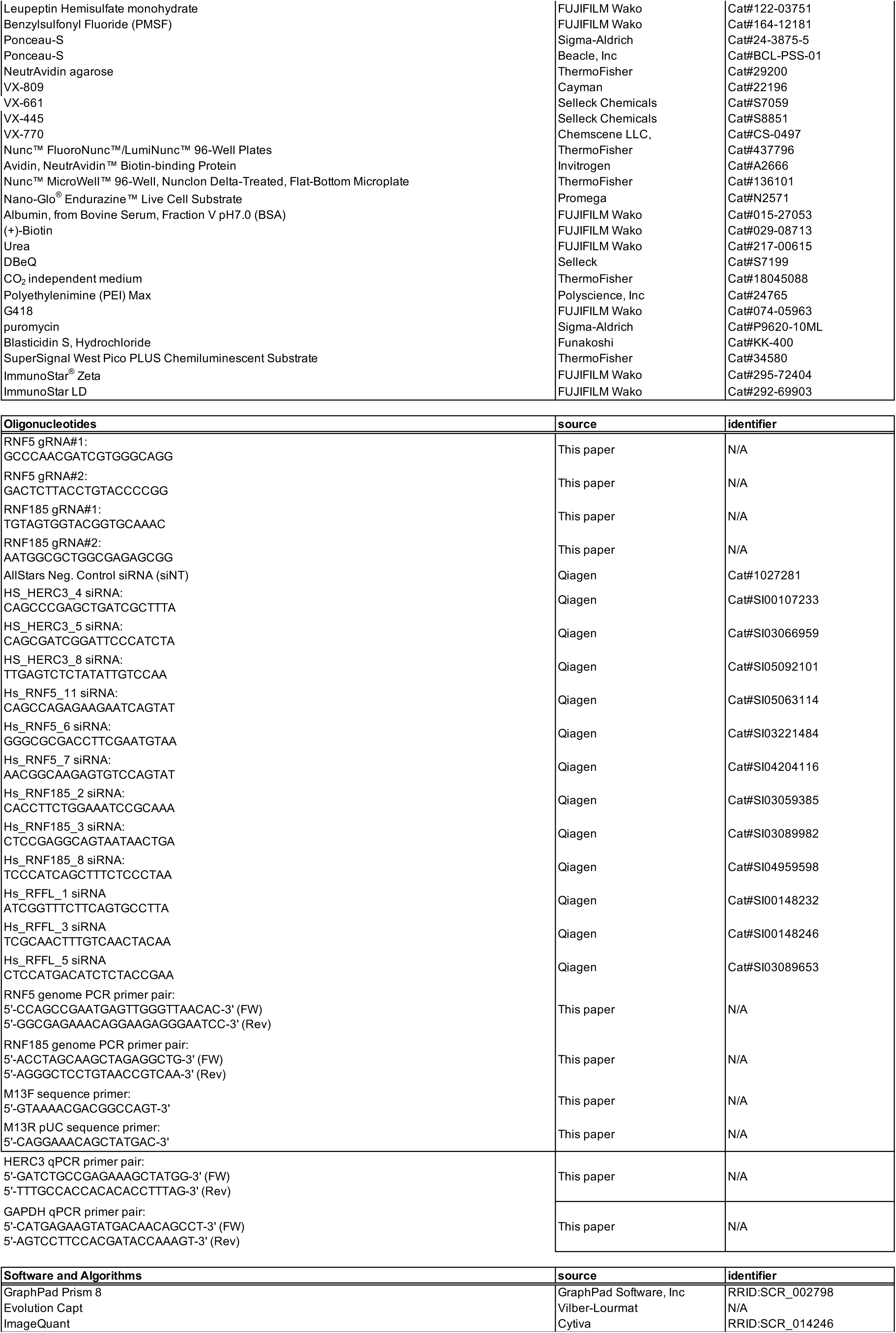

### RESOURCE AVAILABILITY

#### Lead contact

Further information and requests for reagents may be directed to and will be fulfilled by the Lead Contact, Tsukasa Okiyoneda.

#### Materials Availability

Reagents generated in this study are available from the lead contact with a completed materials transfer agreement.

#### Data and Code Availability

All data generated and analyzed during this study are included in this manuscript and the Supplementary Data.

This paper does not report the original code.

Any additional information required to reanalyze the data reported in this paper is available from the lead contact upon request.

## Materials and Methods

### Cell lines and cell culture

CFBE41o-, BHK cells and corresponding transfectants, and COS-7 cells have been previously described (Okiyoneda et al., 2018). 293MSR and RNF5/185 DKO cells were cultured DMEM medium (FUJIFILM Wako Pure Chemical Corporation) supplemented with 10% FBS, 0.5 mg/ml G418. 293MSR and RNF5/185 DKO cells stably expressing HBH-ΔF508-CFTR-3HA were established by lentivirus transduction as previously (Okiyoneda et al., 2018) and were cultured in DMEM medium supplemented with 10% FBS, 0.5 mg/ml G418, and 5 µg/ml blasticidin S. BEAS-2B, 293MSR and CFBE cells stably expressing inducible ΔF508-CFTR-3HA-Nluc were established by lentivirus transduction as previously (Taniguchi et al., 2022). The CFTR expressions in CFBE and BEAS-2B cells were induced by treating them with 1 µg/ml of doxycycline (Dox) for 4 days and 2 days, respectively, unless otherwise specified.

### Plasmids and antibodies

Plasmids and primers used in this study are listed in the key resources table. Truncated, deletion, or mutated constructs of HERC3, UBQLN2, and CFTR were created by PCR-based cloning methods. The HiBiT tag on CFTR variants, Insig-1 and ABCB1 was inserted by PCR-based cloning methods. In most instances, the HiBiT tag was attached to the C-terminus of proteins, situated within the cytosol. For the CFTR-HiBiT(Ex) construct, the extracellular HiBiT(Ex) tag was inserted into the fourth extracellular loop of CFTR, replacing the 3xHA tag on the CFTR-3HA (Okiyoneda et al., 2010). TCRα-HiBiT was created by inserting TCRα from pCMV-TCRα-HA (a kind gift from Dr. R. Kopito) into the pBiT2.1-HiBiT vector by In-fusion cloning. To produce lentivirus vectors, CFTR variants or UBQLN2 variants were first cloned into pDONR221 (Addgene #3388) using BP reaction with BP clonase II Enzyme mix (ThermoFisher). Subsequently, they were transferred to pLX304 (Addgene #25890) or pLIX402 Nluc DEST vector via LR reaction with LR clonase II Enzyme mix (ThermoFisher). The pLIX402 Nluc DEST vector was created by replacing the HA tag in pLIX402 DEST vector (Addgene #41394) with Nluc cDNA in-frame using In-fusion cloning. The pLIX402 ΔF508-CFTR-Nluc (CT) was constructed by LR reaction. CNXss-LgBiT-KDEL was constructed by introducing the N-terminal signal sequence of CNX and the C-terminal KDEL ER retention signal into LgBiT in pBiT1.1-N [TK-LgBiT] (Promega) through PCR-based mutagenesis. Other mutant and chimera constructs were generated through PCR-based mutagenesis or In-fusion cloning.

CRISPR–Cas9 mediated gene knockout of RNF5 or RNF185 was performed using the pSpCas9(BB)-2A-Puro (PX459) V2.0 (Addgene #62988) containing RNF5 gRNA#1 (GCCCAACGATCGTGGGCAGG), RNF5 gRNA#2 (GACTCTTACCTGTACCCCGG), RNF185 gRNA#1 (TGTAGTGGTACGGTGCAAAC), RNF185 gRNA#2 (AATGGCGCTGGCGAGAGCGG). Pairs of oligonucleotides with BpiI (BbsI, ThermoFisher, Cat#ER1012) overhangs were annealed and ligated into the BpiI-digested vector. All constructs generated were verified by DNA sequencing.

The RNF185 antiserum was produced by immunizing a rabbit with recombinant human RNF185 ΔTM (1-130 amino acid) protein purified from E. Coli, and its specificity was confirmed by ELISA. Other antibodies used in this study were listed in the Key Resource Table.

### Mammalian cell transfection

Transient transfection in COS-7, BHK, 293MSR, and RNF5/185 DKO cells was achieved using polyethylenimine (PEI) max (Polysciences Inc), and experiments were performed 2 days post-transfection. siRNA transfections (50 nM each) were performed using Lipofectamine RNAiMax transfection reagent (Invitrogen) according to the manufacturer’s instructions. Cells transfected with siRNA were generally used for the experiments 4-5 days post-transfection unless specified otherwise. The target sequences of siRNA are listed in the Key Resource Table. When not specifically mentioned, pooled siRNA was used for KD. The pooled siRNA was prepared by mixing an equal amount of individual siRNA for HERC3 (siHERC3 #4, #5, #8), RNF5 (siRNF5 #11, #6, #7), RNF185 (siRNF185 #2, #3, #8), and RFFL (siRFFL #1, #3, #5). AllStars Negative Control siRNA from Qiagen was utilized as a negative control non-targeting siRNA (siNT).

### Generation of CRISPR/Cas9-mediated knockout cells

For CRISPR/Cas9-mediated knockouts, 293MSR cells were transfected with specific sgRNAs targeting RNF5 (gRNA#1 and gRNA#2) and RNF185 (gRNA #1 and gRNA #2) using PEI Max (Polyscience, Inc). On the next day, the cells were treated with puromycin (3 µg/mL; Sigma-Aldrich) for 24 hours to enrich the transfected cells. The enriched cells were then diluted and seeded into 96-well plates to allow for the growth of single colonies. Colonies were screened by genome PCR using KOD polymerase (Toyobo) and specific primers listed in the Key Resource Table. The targeted genome deletions were confirmed by genomic DNA sequencing (Figures S1A and S1B) and at the protein level by Western blotting. Confirmation of RNF5 and RNF185 KO was performed through Western blotting and DNA sequencing. For sequencing of RNF5 and RNF185, the genomic locus was amplified by PCR using specific RNF5 and RNF185 genome PCR primer pairs listed in the Key Resource Table. The PCR product was cloned into pMD20-T vector using a Mighty TA-cloning Kit (Takara Bio). The DNA sequence of eight colonies was determined by DNA sequencing using M13F and M13R pUC sequence primers, which resulted in the same result (Figures S1A and S1B).

#### Measurement of PM level and stability of CFTR

Cell-surface expression of ΔF508-CFTR-HRP in CFBE Teton cells on 96 well plates was measured after the addition of the HRP substrate (SuperSignal West Pico PLUS, ThermoFisher) as previously (Okiyoneda et al., 2018; Taniguchi et al., 2022). The subconfluent CFBE cells were transfected with siRNA (50 nM each) in 96-well, and the CFTR expression was induced by 1 µg/ml Dox treatment for 4 days. The cell surface stability of ΔF508-CFTR-3HA in the CFBE cells on 96-well plates was measured by cell surface ELISA using an anti-HA antibody (16B12, BioLegend) as previously (Okiyoneda et al., 2013). Cell surface expression of ΔF508-CFTR was induced by 26°C incubation for 2 days, followed by a 1-hour incubation at 37°C to induce the thermal unfolding. The cellular level and stability of ΔF508-CFTR-3HA in CFBE cells on 96-well plates were measured by ELISA using an anti-HA antibody (16B12, BioLegend) after 4% paraformaldehyde (PFA) fixation and cell permeabilization using 0.1% Triton-X100. After blocking with 0.5% bovine serum albumin (BSA)-PBS, cells were incubated with HRP-conjugated anti-mouse IgG, and the luminescent signal was detected using the HRP substrate (SuperSignal West Pico PLUS, ThermoFisher). The luminescent signal was measured using either the Luminoskan (ThermoFisher), Varioskan Flash microplate reader (ThermoFisher), or the EnSpire Alpha plate reader (Perkin Elmer).

#### Quantitative Real-Time PCR

Total RNA was extracted from cells using TRIzol^®^ (ThermoFisher, Waltham, MA, USA) according to the manufacturer’s protocols. An amount of 500 ng of total RNA was then used for the reverse transcription reaction using ReverTra Ace^®^ qPCR RT Master Mix (Toyobo, Japan). Quantitative RT-PCR was performed in the LightCycler^®^ 480 System (Roche Diagnostics, Switzerland), and the gene expression was examined by SYBR Advantage qPCR Premix (Toyobo, Japan). The relative quantity of the target gene mRNA was normalized using human GAPDH as the internal control. PCR amplification was performed in 2 steps, and the reaction protocol included preincubation at 95˚C for 3 min; then the amplification of 40 cycles was set for 5 s at 95˚C and 30 s at 60˚C, and the melting curve of 5 min at 95˚C, 60 s at 60˚C, and 97˚C. The sequences of primers used for quantitative RT-PCR are listed in the Key Resource Table.

#### Halide-Sensitive YFP Quenching Assay

ΔF508-CFTR function assay by halide-sensitive YFP fluorescence quenching was performed as described (Okiyoneda et al., 2018; Veit et al., 2014). CFBE cells expressing both inducible ΔF508-CFTR-3HA and halide sensor YFP-F46L/H148Q/I152L were seeded onto black 96-well microplates and transfected with siRNA (50 nM each). The CFTR expression was induced by 1 µg/ml Dox treatment for 4 days. Cell surface expression of low temperature-rescued ΔF508-CFTR was induced for 2 days at 26°C, followed by a 1-hour incubation at 37°C before analysis. At the time of assay, cells were washed four times with 400 µL of phosphate-buffered saline (PBS)-chloride (140 mM NaCl, 2.7 mM KCl, 8.1 mM Na_2_HPO_4_, 1.5 mM KH_2_PO_4_, 1.1 mM MgCl_2_, 0.7 mM CaCl_2_, and 5 mM glucose) and incubated with PBS-chloride (50 µL per well), followed by well wise-injection of an activator solution (50 µL per well) containing 20 µM forskolin (FUJIFILM Wako Pure Chemical Corporation), 0.5 mM 3-isobutyl-1-methyl-xanthine (IBMX, FUJIFILM Wako Pure Chemical Corporation), 0.5 mM 8-(4-chlorophenylthio)-adenosine-3’,5’-cyclic monophosphate (CPT-cAMP, Santa Cruz Biotechnology), and 0.1 mM genistein (FUJIFILM Wako Pure Chemical Corporation)]. The cells were then incubated for 57 seconds. The fluorescence was continuously recorded (200 ms per point) for 3 seconds (baseline) and for 32 seconds after the rapid addition of 100 µL PBS-iodide, where NaCl was replaced with NaI. The fluorescence measurements were carried out using a VICTOR Nivo multimode microplate reader (Perkin Elmer) with a dual syringe pump (excitation/emission 500/535 nm). After normalizing the YFP signals before PBS-iodide injection, the I^-^ influx rate was calculated by curve-fitting the YFP fluorescence decay using GraphPad Prism 8 software (GraphPad Software).

#### Western blotting

Cells were solubilized in a RIPA buffer supplemented with 1 mM PMSF (FUJIFILM Wako Pure Chemical Corporation), 5 µg/ml leupeptin (FUJIFILM Wako Pure Chemical Corporation), and 5 µg/ml pepstatin A (Peptide Institute. INC.). The cell lysates were analyzed by a Western blot as previously described (Okiyoneda et al., 2010). Western blots were visualized using a SuperSignal West Pico PLUS Chemiluminescent Substrate (ThermoFisher), ImmunoStar Zeta (FUJIFILM Wako Pure Chemical Corporation), or ImmunoStar LD (FUJIFILM Wako Pure Chemical Corporation) and analyzed by FUSION Chemiluminescence Imaging System (Vilber Bio Imaging). In some cases, the staining of Ponceau S was used as a loading control.

#### CFTR Ubiquitination Measurement by Western blotting

CFTR ubiquitination was measured as previously (Okiyoneda et al., 2010; Okiyoneda et al., 2018). Briefly, CFBE Teton HBH-ΔF508-CFTR-3HA cells, 293MSR, and RNF5/185 DKO cells were lysed in RIPA buffer containing 5 µg/mL Leupeptin, 5 µg/mL Pepstatin A, 100 µM PMSF, 10 µM MG-132, and 5 mM N-ethylmaleimide (NEM) after 10 µM MG-132 treatment at 37°C for 1 hour (CFBE) or 3 hours (293MSR, RNF5/185 DKO). The CFTR expression was induced by 1 µg/ml Dox treatment for 4 days. Immature HBH-ΔF508-CFTR was purified using Neutravidin agarose (ThermoFisher) under denaturing conditions and analyzed by Western blotting with anti-Ub (P4D1) or anti-K48 (Apu2) and anti-HA antibodies.

#### Ub ELISA

CFTR ubiquitination levels in CFBE cells, 293MSR, and RNF5/185 DKO cells were performed as previously (Kamada et al., 2019; Okiyoneda et al., 2018). Cells were lysed in RIPA buffer supplemented with 5 µg/mL Leupeptin, 5 µg/mL Pepstatin A, 100 µM PMSF, 10 µM MG-132, and 5 mM NEM after 10 µM MG-132 treatment at 37°C for 1 hour (CFBE) or 3 hours (293MSR, RNF5/185 DKO). The HBH-ΔF508-CFTR in the cell lysate was immobilized on NA-coated 96 well-white plates and denatured in 8 M urea at room temperature for 5 min. After the denaturation, the plate was washed 5 times with 2 M Urea in RIPA buffer. Following 4 washes with 0.1% NP-40-PBS, the plate was blocked with 0.1% BSA. CFTR ubiquitination was detected using anti-K48 Ub (Apu2, Merck Millipore) and anti-K63 Ub (Apu3, Merck Millipore) antibodies. The ubiquitination levels were quantified with an HRP-conjugated secondary antibody. Linkage-specific CFTR ubiquitination levels were normalized for the CFTR level quantified using the anti-HA antibody (16B12, BioLegend).

#### Pull-down experiments

To detect the interaction of HBH-ΔF508-CFTR-3HA and Myc-HERC3 and/or FLAG-UBQLN2, BHK cells stably expressing HBH-ΔF508-CFTR-3HA were transfected with Myc-HERC3 and/or FLAG-UBQLN2. Two days post-transfection, the cells were solubilized in mild lysis buffer (150 mM NaCl, 20 mM Tris, 0.1% NP-40, pH 8.0) supplemented with 1 mM PMSF, 5 mg/ml leupeptin, and pepstatin. The cell lysates were then incubated with NA agarose (ThermoFisher) for 2 hours at 4˚C. After being washed 4 times with mild lysis buffer, the complex was eluted in urea elution buffer (8 M urea, 2% SDS, 3 mM biotin) at room temperature for 30 min and analyzed by Western blotting. In Figures 4A and 5G, cells were treated with 10 µM MG-132 for 3 hours before cell lysis.

#### HiBiT and Nluc degradation assay

For the HiBiT degradation assay, subconfluent 293MSR and RNF5/185 DKO cells in 6-well plates were transfected with siRNA (50 nM each). After one day of culture, the cells were trypsinized, reseeded in 6-well plates, and cultured for another day. Then, the cells were transfected with different HiBiT fusion constructs, including CFTR variants-HiBiT (CT), TCRα-HiBiT, Insig-1-HiBiT, or ΔY490-ABCB1-HiBiT, along with cytosolic LgBiT (pBiT1.1-N [TK/LgBiT], Promega). The following day, the cells were trypsinized again and seeded in Nunc™ MicroWell™ 96-Well Nunclon Delta-Treated Flat-Bottom Microplates (ThermoFisher) and cultured for 18-24 hours.

For the Nluc degradation assay, subconfluent BEAS-2B or CFBE cells stably expressing Dox-inducible ΔF508-CFTR-Nluc were transfected with siRNA (50 nM each) in 6-well plates. The next day, the cells were trypsinized and seeded in 96-well plates (2 x 10^4^ cells/well) and cultured for 18-24 hours. The expression of ΔF508-CFTR-Nluc was induced by treating the cells with 1 µg/mL Dox for 2 days. After washing the cells with 100 µL of CO_2_-independent medium (ThermoFisher), 50 µL of 0.1x Nano-Glo^®^ Endurazine (Promega) in CO_2_-independent medium was added to each well, and the cells were incubated for 2.5 hours at 37˚C in 5% CO_2_. To measure the degradation kinetics, 10 µL of 600 µg/mL CHX was added to each well, resulting in a final concentration of 100 µg/mL. Luminescence was continuously measured at 5-minute intervals for 3-12 hours using a Luminoskan plate reader (ThermoFisher). The luminescence signal of CHX-treated cells (Lum^CHX^) was normalized to the signal of untreated cells (Lum^untreat^) to calculate the remaining ERAD substrates during the CHX chase as equation (1). The ERAD rate was calculated by fitting the luminescence decay with a one-phase exponential decay function using GraphPad Prism 8 (GraphPad Software, La Jolla, CA, USA).

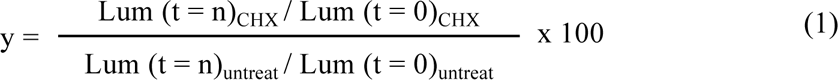

### HiBiT retro-translocation assay

Subconfluent cells were transfected with ΔF508-CFTR-HiBiT (Ex) and cytosolic LgBiT (pBiT1.1-N [TK/LgBiT], Promega). The next day, cells were trypsinized and seeded in 96-well plates (2 x10^4^ cells/well) and cultured for 18-24 hours. Cells were washed with 100 µL of the CO_2_-independent medium. Then, 50 µL of 0.5x Nano-Glo^®^ Endurazine (Promega) in the CO_2_-independent medium was added to each well. The cells were incubated for 2.5 hours at 37°C and 5% CO_2_. To detect the retro-translocated ΔF508-CFTR signal, 10 µL of 60 µM MG-132 (final 10 µM) with or without 600 µg/ml CHX (final 100 µg/ml) in the CO_2_-independent medium was added to each well. Luminescence was continuously measured at 5-min intervals for 3 hours using a Luminoskan plate reader (ThermoFisher). The amount of retrotranslocated CFTR was calculated by subtracting the luminescence signal before MG-132 treatment from the signal after MG-132 treatment. The retrotranslocation rate was then calculated by linear fitting the increased retrotranslocation during MG-132 treatment. In the Trikafta experiments, cells were incubated with Trikafta (consisting of 3 µM VX-661, 3 µM VX-445, and 1 µM VX-770) for 24 hours at 37°C. Trikafta treatment was also maintained during Endurazine loading and the MG-132 chase phase when luminescent measurements were conducted.

#### HiBiT ER disappearance assay

Subconfluent cells were transfected with ΔF508-CFTR-HiBiT (Ex) and ER luminal LgBiT (CNXss-LgBiT-KDEL). The CNXss-LgBiT-KDEL construct contains a signal sequence of calnexin (CNX) fused with LgBiT at the N-terminus and an ER retention signal (KDEL) fused with LgBiT at the C-terminus. The next day, the transfected cells were trypsinized and seeded in 96-well plates (2 x10^4^ cells/well) and cultured for 18-24 hours. After washing cells with 100 µL of the CO_2_-independent medium, 50 µL of 0.1x Nano-Glo^®^ Endurazine^TM^ (Promega) in the CO_2_-independent medium was added to each well. The cells were then incubated for 2.5 hours at 37˚C, 5% CO_2_. To measure the ER disappearance of CFTR-HiBiT(Ex), 10 µL of 600 µg/mL CHX in the CO_2-_independent medium was added to each well (final 100 µg/ml). Luminescence was continuously measured at 5-min intervals for 3 hours using a Luminoskan plate reader (ThermoFisher). The luminescence signal (Lum) of CHX-treated cells (Lum^CHX^) was normalized to the signal of untreated cells (Lum^untreat^) to calculate the decrease in luminescence signal due to CHX treatment as equation (1) like the HiBiT and Nluc degradation assay. The ERAD rate was calculated by fitting the luminescence decay with a one-phase exponential decay function using GraphPad Prism 8

#### ELISA to detect CFTR-UBQLN2 interaction

293MSR cells co-transfected with HBH-ΔF508-CFTR-3HA and FLAG-UBQLN2 were treated with 10 µM MG-132 for 1 hour at 37˚C and incubated with 0.1% PFA in PBS (+) for 15 min for crosslinking nearby proteins. After quenching by adding 125 mM glycine, cells were washed with PBS (+) twice and lysed in RIPA buffer containing 5 µg/mL leupeptin, 5 µg/mL pepstatin A, and 100 µM PMSF. HBH-ΔF508-CFTR-3HA in cell lysate was immobilized on an NA-coated 96-plate at 4 ˚C for 2 hours. After 5 times washing with 0.1% NP-40 in PBS (-) and blocking with 0.5% BSA in 0.1% NP-40 PBS (-), HBH-ΔF508-CFTR and FLAG-UBQLN2 were probed with anti-HA (16B12) and anti-FLAG (1E6) antibodies, respectively, and quantified with an HRP-conjugated secondary antibody. After incubating with SuperSignal West Pico PLUS Chemiluminescent Substrate (ThermoFisher), the luminescent signals were measured using a Variokan plate reader (ThermoFisher). The specific signals of Flag-UBQLN2 and HBH-ΔF508-CFTR-3HA were quantified by subtracting the background signals obtained from the lysates of cells transfected with FLAG-UBQLN2 alone, without HBH-ΔF508-CFTR-3HA. FLAG-UBQLN2 binding on the HBH-ΔF508-CFTR-3HA was normalized for the CFTR level quantified by an anti-HA antibody (16B12).

### Subcellular fractionation

Microsomes were isolated from 293MSR cells stably expressing HBH-ΔF508-CFTR-3HA treated with or without 10 µM MG-132 for 3 hours at 37˚C. In brief, cells were resuspended with resuspension buffer (10 mM HEPES, pH 7.5, 0.25 M Sucrose, 10 mM KCl, 1.5 mM MgCl_2_, 1 mM EDTA, 1 mM EGTA, 5 µg/mL leupeptin, 5 µg/mL pepstatin A, and 100 µM PMSF), and sheared by passing 30 times with 25G needle. The cell homogenates were centrifuged at 1,000 g at 4˚C for 5 min, and supernatants recentrifuged 100,000 g at 4˚C for 1 hour. The supernatant was used as cytosol, and the pellet fraction containing microsome was dissolved with RIPA buffer. Both microsome and cytosol were analyzed with Western blotting and the UBQLN2 abundance was quantified by densitometry. The ER recruited UBQLN2 was calculated as equation (2) and normalized by the abundance of recruited UBQLN in the WT cells transfected with siNT.

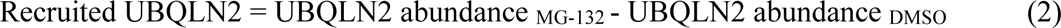

### DSP Cross-link Pull-Down Assay

293MSR WT and RNF5/185 DKO cells stably expressing HBH-ΔF508-CFTR-3HA were transfected with siRNA (50 nM each) and treated with 10 µM MG-132 for 1 hour at 37˚C. After washing with PBS(+), cells were incubated with 0.1 mM dithiobis[succinimidyl propionate] (DSP, ThermoFisher) in PBS(+) for 15 min at room temperature for cross-linking. After washing with PBS(+) and 0.5% BSA in PBS(+) several times, cells were solubilized in mild lysis buffer (150 mM NaCl, 20 mM Tris, 0.1% NP-40, pH 8.0, supplemented with 1 mM PMSF, 5 mg/ml leupeptin and pepstatin). Cell lysates were incubated with Neutravidin agarose (ThermoFisher) for 2 hours at 4˚C. After being washed 5 times with mild lysis buffer, the complex was eluted in urea elution buffer (8 M urea, 2% SDS, 3 mM biotin) at 30˚C for 30 min and analyzed by Western blotting.

#### Statistical analysis

For quantification, data from at least 3 independent experiments were used where the data are expressed as means ± standard deviation (SD). Statistical significance was assessed from at least three biological replicates by one-way or two-way ANOVA or Student’s t-test as specified in figure legends using GraphPad Prism 8. A P value < 0.05 was defined as statistically significant.

