## Supplemental Information for "HERC3 E3 ligase provides an ERAD branch eliminating select membrane proteins"

Running title: The novel ERAD branch mediated by cytosolic HERC3

Yuka Kamada[1], Yuko Ohnishi[1], Chikako Nakashima[1], Aika Fujii[1], Mana Terakawa[1], Ikuto Hamano[1], Uta Nakayamada[1], Saori Katoh[1], Noriaki Hirata[1], Hazuki Tateishi[1], Ryosuke Fukuda[1], Hirotaka Takahashi[2], Gergely L. Lukacs[3,4], Tsukasa Okiyoneda[1,5]

[1]Department of Biomedical Sciences, School of Biological and Environmental Sciences, Kwansei Gakuin University, Sanda 669-1330, Hyogo, Japan.
[2]Division of Cell-Free Sciences, Proteo-Science Center (PROS), Ehime University, Matsuyama 790-8577, Ehime, Japan.
[3]Department of Physiology, [4]Department of Biochemistry, McGill University, Montréal, QC H3G 1Y6, Canada.

[5]corresponding:

**Supplemental Information**

**Figure S1.**

**Establishment of RNF5/185 DKO 293MSR cells.**

(A, B) Schematic representation of the RNF5 (A) and RNF185 (B) -targeting gRNA sequences. Arrows indicate primer positions. PAM, protospacer adjacent motif. The locations of each start codon, stop codon and the catalytic cysteine residues of RNF5 (C42) and RNF185 (C39, C42) are also indicated. The sequences analyzed for each KO cell line are shown along with the deleted sequences.

**Figure S2.**

**A representative luminescence trace from the ΔF508-CFTR-HiBiT(Ex) retrotranslocation assay**

293MSR cells were transiently transfected with LgBiT, with or without ΔF508-CFTR-HiBiT(Ex), followed by Endurazine loading. Luminescence was continuously monitored in live cells during treatment with or without 10 μM MG-132. This figure is associated with Figure 4B.

**Figure S3.**

**The correlation analysis**

(A) The relationship between CFTR ERAD (Fig.3F) and the retrotranslocation rates of ΔF508-CFTR (Fig.4D) was analyzed for correlation.

(B) Correlation analysis was performed to examine the connection between CFTR ER disappearance (Fig.4F) and the retrotranslocation rates of ΔF508-CFTR (Fig. 4D).

(C) The correlation between CFTR K48-linked polyubiquitination (Fig.5B) and K63-linked polyubiquitination (Fig.5C).

(D) The correlation between CFTR K48-linked polyubiquitination (Fig.5B) and the ERAD rate (Fig.3F).

(E) The correlation between CFTR K63-linked polyubiquitination (Fig.5C) and the ERAD rate (Fig.3F).

(F) The correlation between CFTR K48-linked polyubiquitination (Fig.5B) and the retrotranslocation rate of ΔF508-CFTR (Fig.4D).

(G) The correlation between CFTR K63-linked polyubiquitination (Fig.5C) and the retrotranslocation rate of ΔF508-CFTR (Fig.4D).

(H) The correlation between the CFTR K48-linked polyubiquitination (Fig.5B) and UBQLN2 binding (Fig.6C).

(I) The correlation between the CFTR K63-linked polyubiquitination (Fig.5C) and UBQLN2 binding (Fig.6C).
(J) The correlation between the CFTR ERAD (Fig.3F) and UBQLN2 binding (Fig.6C).
(K) The correlation between the retrotranslocation (Fig.4D) and UBQLN2 binding (Fig.6C).

**Figure S4.**

**Effects of UBQLN single KD on the CFTR ERAD, triple KD on CFTR retrotranslocation, and establishment of TCRα-HiBiT and Insig-1-HiBiT ERAD assay.**

(A) The kinetic degradation of ΔF508-CFTR-HiBiT(CT) in 293MSR WT cells transfected with 50 nM siNT or siUNQLN1, 2, or 4. The ERAD rate was calculated by fitting the initial degradation portion of each kinetic degradation curve (right, n=2).
Each biological replicate (n) is color-coded: the averages from 4 technical replicates are shown in triangles. Data represent mean.
(B) The retrotranslocation of ΔF508-CFTR-HiBiT(Ex) in 293MSR cells upon UBQLN1/2/4 triple KD was measured during the MG-132 and CHX chase (n=3, unpaired t-test).
(C, D) The HiBiT degradation assay confirmed the proteasomal degradation of TCRα-HiBiT (C, n=4) and Insig-1-HiBiT (D, n=3) in 293MSR cells.
Data represent mean ± SD. **p < 0.01.

**Figure S5.**

**Effects of ablation of HERC3 and/or RNF5/185 on the ERAD of ΔF508-NBD1 and N1303K-CFTR, correlation of the ERAD between ΔF508-CFTR and CFTR fragments.**

(A, B) The HiBiT degradation assay measured the ERAD of ΔF508-NBD1-HiBiT (A, n=4) and N1303K-CFTR-HiBiT (B, n=3) in 293MSR WT and RNF5/185 KO cells transfected with 50 nM siNT or siHERC3 as indicated. Two-way RM ANOVA revealed a significant main effect of HERC3 KD (A, B) or RNF5/185 DKO (B), but no interaction between them ($P_{int}$>0.05, in A, B). Each biological replicate (n) is color-coded: the averages from 4 technical replicates are shown in triangles (A, B). Data represent mean ± SD. *p < 0.05, **p < 0.01, ****p < 0.0001, ns, not significant.
(C, D, E) The correlation of ERAD rates between ΔF508-CFTR (Fig.3F) and M1 (C, Fig.9B), M1-N1(ΔF) (D, Fig.9C), or M2 (E, Fig.9D).

### Figure S1

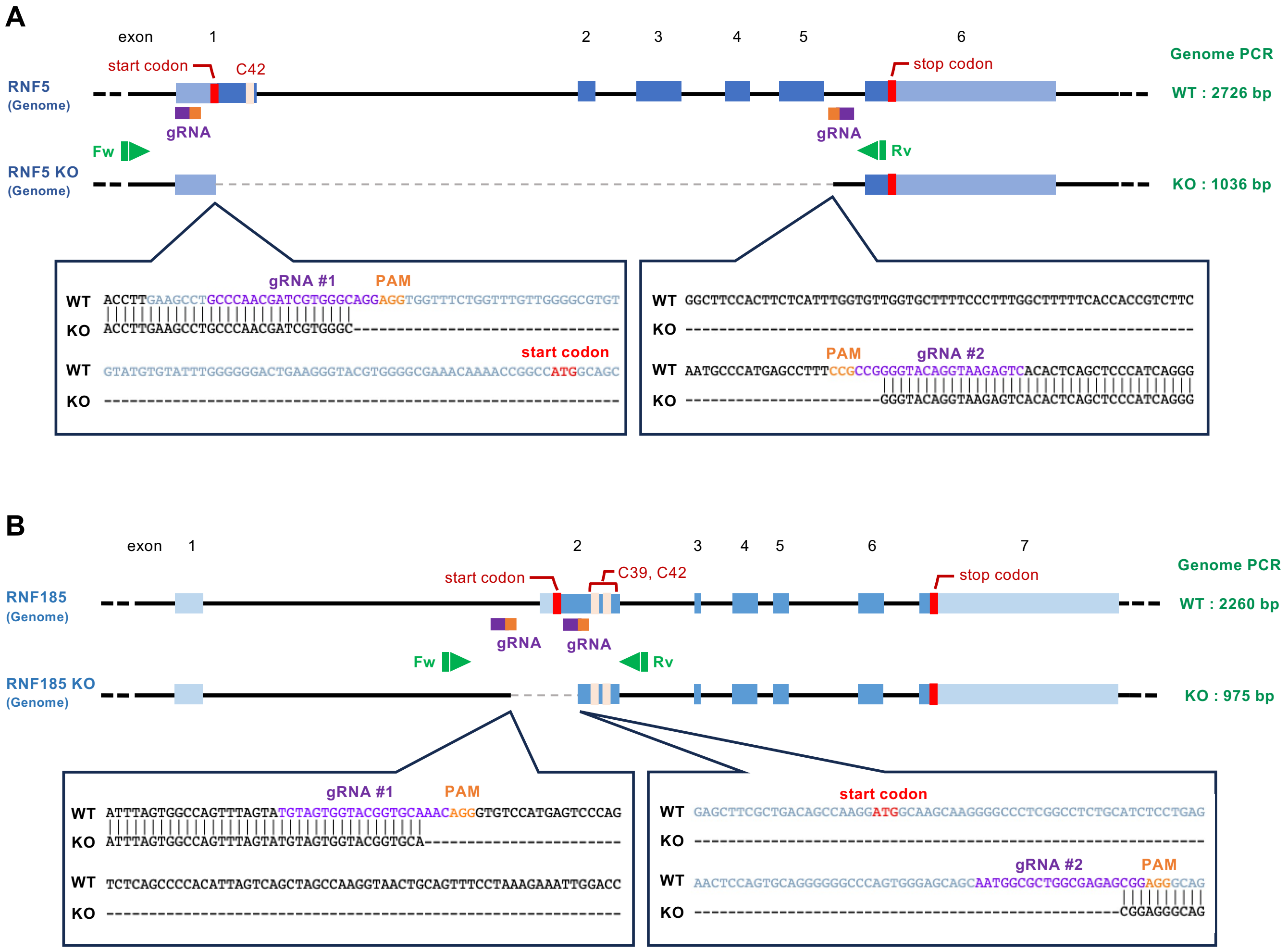
A
exon 1 2 3 4 5 6
RNF5
(Genome)
start codon
C42
stop codon
Genome PCR
WT : 2726 bp
gRNA
gRNA
Fw
Rv
RNF5 KO
(Genome)
KO : 1036 bp
gRNA #1
PAM
WT ACCTTGAAGCCTGCCCAACGATCGTGGGCAGGAGGTGGTTTCTGGTTTGTTGGGGCGTGT
KO ACCTTGAAGCCTGCCCAACGATCGTGGGC--------------------------------
start codon
WT GTATGTGTATTTGGGGGACTGAAGGGTACGTGGGGCGAAACAAAACCGGCCATGGCAGC
KO -----------------------------------------------------------
WT GGCTTCCACTTCTCATTTGGTGTTGGTGCTTTTCCCTTTGGCTTTTTCACCACCGTCTTC
KO ------------------------------------------------------------
PAM
gRNA #2
WT AATGCCCATGAGCCTTTCCGCCGGGGTACAGGTAAGAGTCACACTCAGCTCCCATCAGGG
KO -----------------------GGGTACAGGTAAGAGTCACACTCAGCTCCCATCAGGG
B
exon 1 2 3 4 5 6 7
RNF185
(Genome)
start codon
C39, C42
stop codon
Genome PCR
WT : 2260 bp
gRNA
gRNA
Fw
Rv
RNF185 KO
(Genome)
KO : 975 bp
gRNA #1
PAM
WT ATTTAGTGGCCAGTTTAGTATGTAGTGGTACGGTGCAAACAGGGTGTCCATGAGTCCCAG
KO ATTTAGTGGCCAGTTTAGTATGTAGTGGTACGGTGCA------------------------
WT TCTCAGCCCCACATTAGTCAGCTAGCCAAGGTAACTGCAGTTTCCTAAAGAAATTGGACC
KO ------------------------------------------------------------
start codon
WT GAGCTTCGCTGACAGCCAAGGATGGCAAGCAAGGGGCCCTCGGCCTCTGCATCTCCTGAG
KO ------------------------------------------------------------
gRNA #2
PAM
WT AACTCCAGTGCAGGGGGGCCCAGTGGGAGCAGCAATGGCGCTGGCGAGAGCGGAGGGCAG
KO --------------------------------------------------CGGAGGGCAG

### Figure S2

**A**

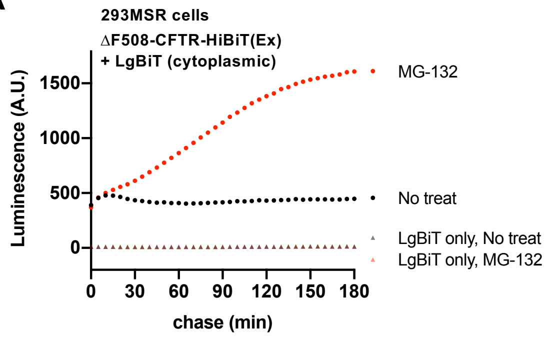

293MSR cells
ΔF508-CFTR-HiBiT(Ex)
+ LgBiT (cytoplasmic)
Luminescence (A.U.)
1500
1000
500
0
0
30
60
90
120
150
180
chase (min)
MG-132
No treat
LgBiT only, No treat
LgBiT only, MG-132

### Figure S3

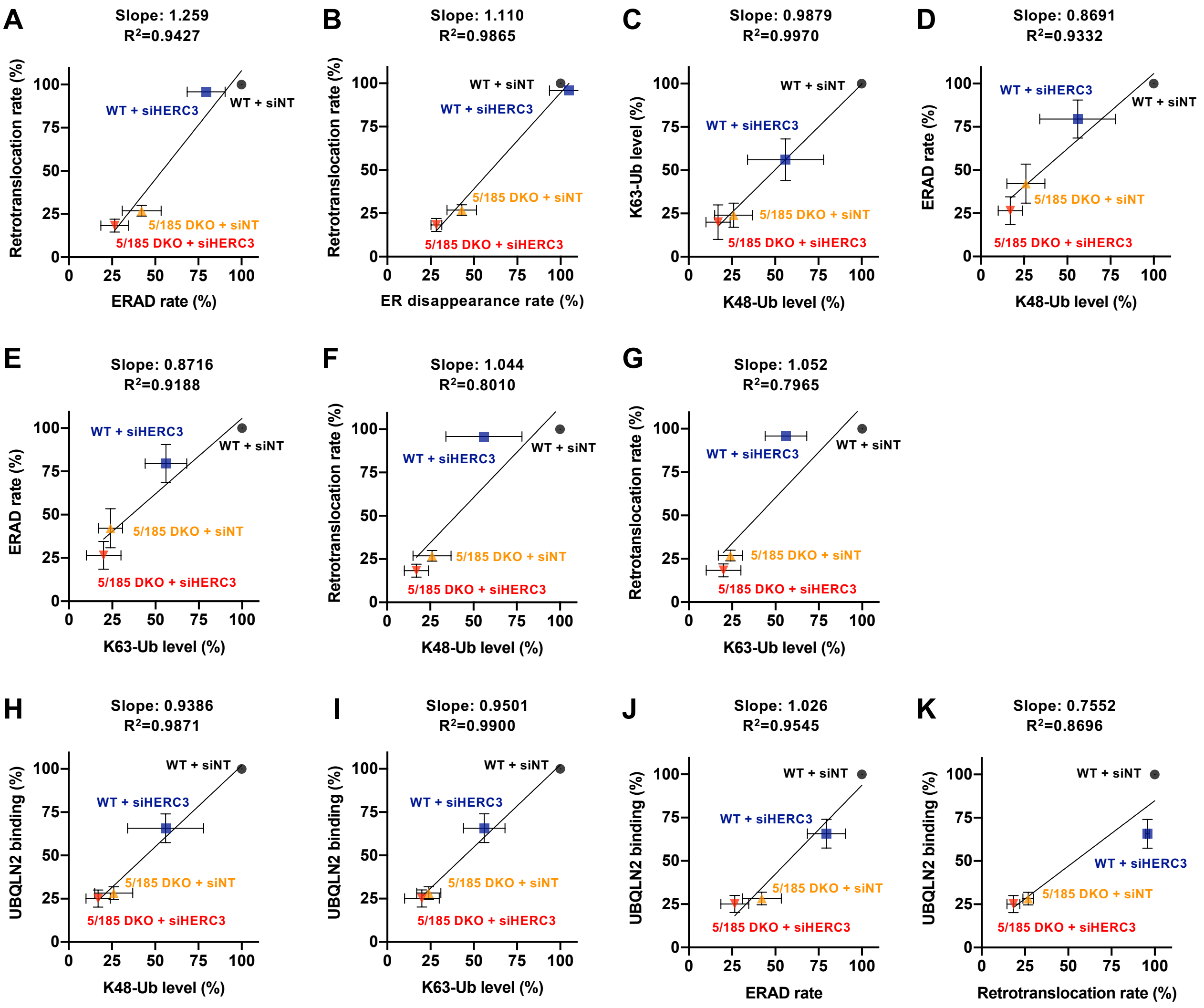
A
Slope: 1.259
R²=0.9427
Retrotranslocation rate (%)
ERAD rate (%)
WT + siHERC3
WT + siNT
5/185 DKO + siNT
5/185 DKO + siHERC3
B
Slope: 1.110
R²=0.9865
Retrotranslocation rate (%)
ER disappearance rate (%)
WT + siNT
WT + siHERC3
5/185 DKO + siNT
5/185 DKO + siHERC3
C
Slope: 0.9879
R²=0.9970
K63-Ub level (%)
K48-Ub level (%)
WT + siNT
WT + siHERC3
5/185 DKO + siNT
5/185 DKO + siHERC3
D
Slope: 0.8691
R²=0.9332
ERAD rate (%)
K48-Ub level (%)
WT + siHERC3
WT + siNT
5/185 DKO + siNT
5/185 DKO + siHERC3
E
Slope: 0.8716
R²=0.9188
ERAD rate (%)
K63-Ub level (%)
WT + siHERC3
WT + siNT
5/185 DKO + siNT
5/185 DKO + siHERC3
F
Slope: 1.044
R²=0.8010
Retrotranslocation rate (%)
K48-Ub level (%)
WT + siHERC3
WT + siNT
5/185 DKO + siNT
5/185 DKO + siHERC3
G
Slope: 1.052
R²=0.7965
Retrotanslocation rate (%)
K63-Ub level (%)
WT + siHERC3
WT + siNT
5/185 DKO + siNT
5/185 DKO + siHERC3
H
Slope: 0.9386
R²=0.9871
UBQLN2 binding (%)
K48-Ub level (%)
WT + siNT
WT + siHERC3
5/185 DKO + siNT
5/185 DKO + siHERC3
I
Slope: 0.9501
R²=0.9900
UBQLN2 binding (%)
K63-Ub level (%)
WT + siNT
WT + siHERC3
5/185 DKO + siNT
5/185 DKO + siHERC3
J
Slope: 1.026
R²=0.9545
UBQLN2 binding (%)
ERAD rate
WT + siNT
WT + siHERC3
5/185 DKO + siNT
5/185 DKO + siHERC3
K
Slope: 0.7552
R²=0.8696
UBQLN2 binding (%)
Retrotranslocation rate (%)
WT + siNT
WT + siHERC3
5/185 DKO + siNT
5/185 DKO + siHERC3

### Figure S4

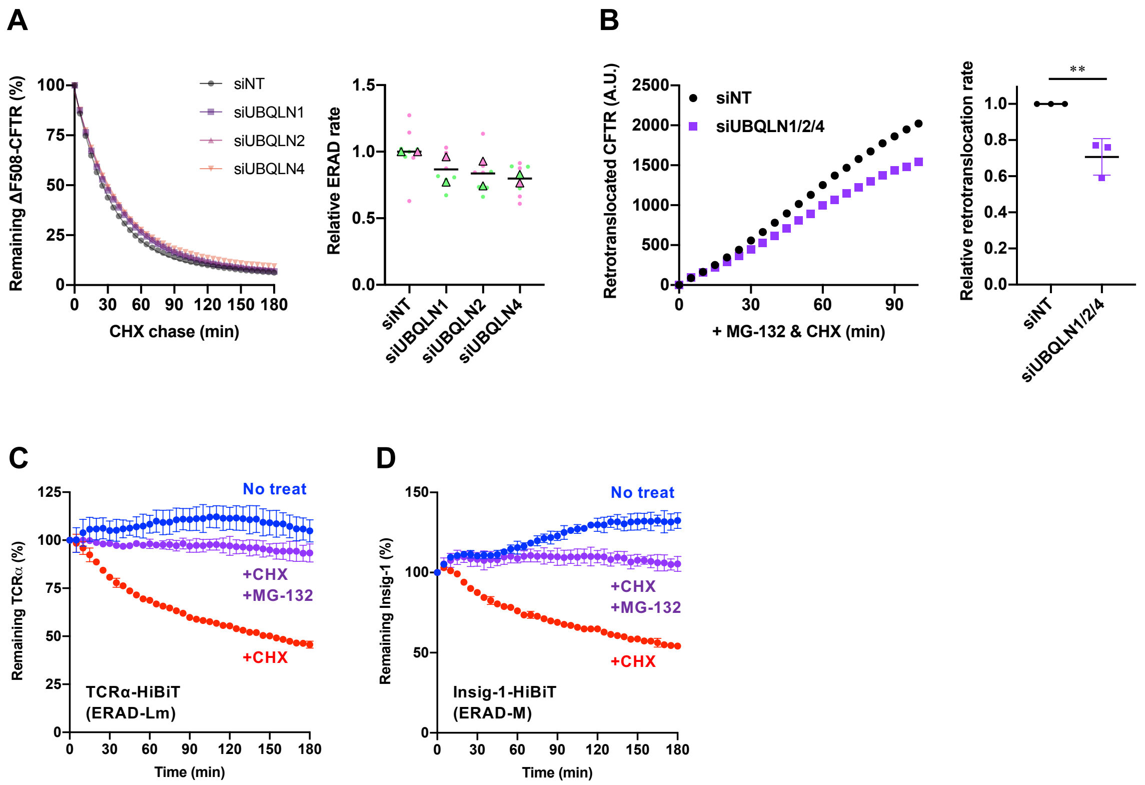

A
Remaining ΔF508-CFTR (%)
CHX chase (min)
siNT
siUBQLN1
siUBQLN2
siUBQLN4
Relative ERAD rate
B
Retrotranslocated CFTR (A.U.)
+ MG-132 & CHX (min)
siNT
siUBQLN1/2/4
Relative retrotranslocation rate
**
C
Remaining TCRα (%)
No treat
+CHX
+MG-132
+CHX
TCRα-HiBiT
(ERAD-Lm)
Time (min)
D
Remaining Insig-1 (%)
No treat
+CHX
+MG-132
+CHX
Insig-1-HiBiT
(ERAD-M)
Time (min)

### Figure S5

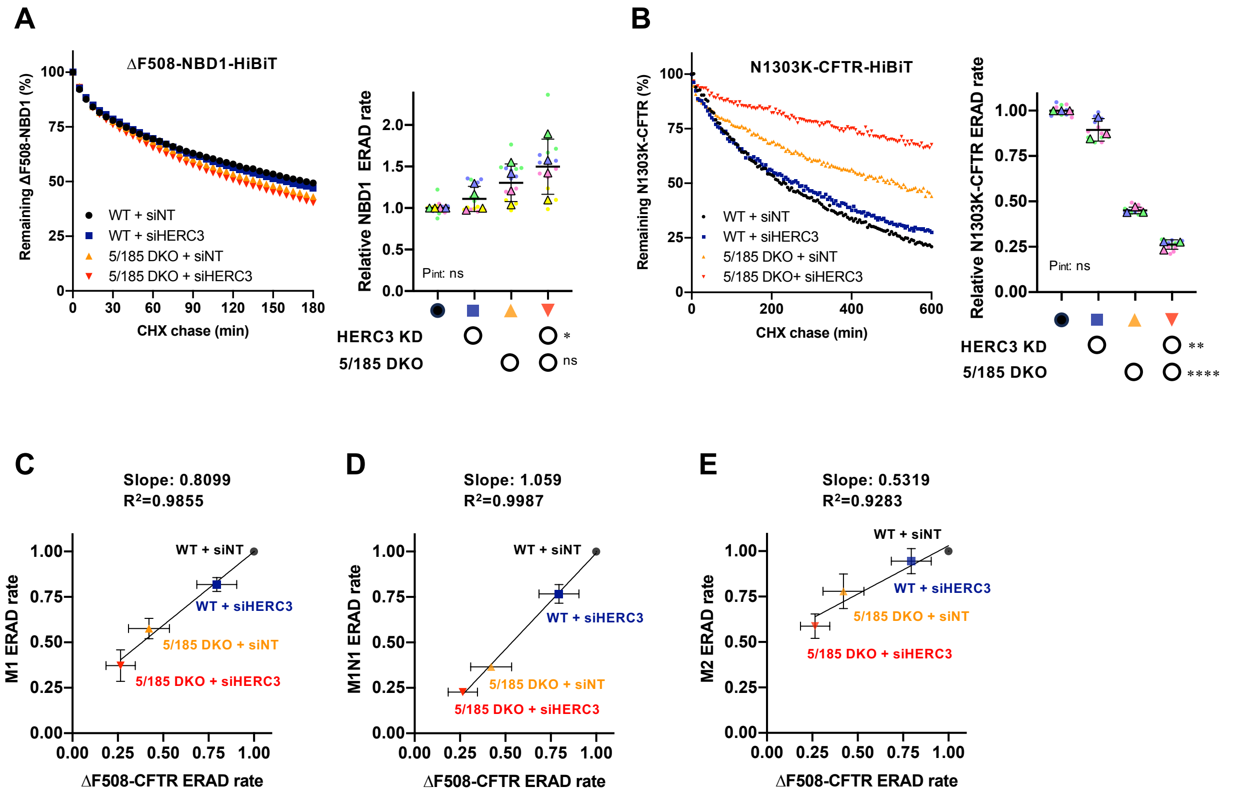

A
ΔF508-NBD1-HiBiT
Remaining ΔF508-NBD1 (%)
CHX chase (min)
WT + siNT
WT + siHERC3
5/185 DKO + siNT
5/185 DKO + siHERC3
Relative NBD1 ERAD rate
Pint: ns
HERC3 KD
5/185 DKO
*
ns
B
N1303K-CFTR-HiBiT
Remaining N1303K-CFTR (%)
CHX chase (min)
WT + siNT
WT + siHERC3
5/185 DKO + siNT
5/185 DKO+ siHERC3
Relative N1303K-CFTR ERAD rate
Pint: ns
HERC3 KD
5/185 DKO
**
****
C
Slope: 0.8099
R²=0.9855
M1 ERAD rate
ΔF508-CFTR ERAD rate
WT + siNT
WT + siHERC3
5/185 DKO + siNT
5/185 DKO + siHERC3
D
Slope: 1.059
R²=0.9987
M1N1 ERAD rate
ΔF508-CFTR ERAD rate
WT + siNT
WT + siHERC3
5/185 DKO + siNT
5/185 DKO + siHERC3
E
Slope: 0.5319
R²=0.9283
M2 ERAD rate
ΔF508-CFTR ERAD rate
WT + siNT
WT + siHERC3
5/185 DKO + siNT
5/185 DKO + siHERC3
